# USP7 cooperates with NOTCH1 to drive the oncogenic transcriptional program in T cell leukemia

**DOI:** 10.1101/248427

**Authors:** Kelly M. Arcipowski, Carlos A. Martinez, Qi Jin, Yixing Zhu, Blanca Teresa Gutierrez Diaz, Kenneth K. Wang, Megan R. Johnson, Andrew G. Volk, Feng Wang, Jian Wu, Hui Wang, Ivan Sokirniy, Paul M. Thomas, Young Ah Goo, Nebiyu A. Abshiru, Nobuko Hijiya, Sofie Peirs, Niels Vandamme, Geert Berx, Steven Goosens, Stacy A. Marshall, Emily J. Randleman, Yoh-hei Takahashi, Lu Wang, Elizabeth T. Bartom, Clayton K. Collings, Pieter Van Vlierberghe, Alexandros Strikoudis, Stephen Kelly, Beatrix Ueberheide, Christine Mantis, Irawati Kandela, Jean-Pierre Bourquin, Beat Bornhauser, Valentina Serafin, Silvia Bresolin, Maddalena Paganin, Benedetta Accordi, Giuseppe Basso, Neil L. Kelleher, Joseph Weinstock, Suresh Kumar, John D. Crispino, Ali Shilatifard, Panagiotis Ntziachristos

## Abstract

T-cell acute lymphoblastic leukemia (T-ALL) is an aggressive disease, affecting children and adults. Treatments^1-6^ show high response rates but have debilitating effects and carry risk of relapse^5,7,8^. Previous work implicated NOTCH1 and other oncogenes^1,2,9-20^. However, direct inhibition of these pathways affects healthy tissues and cancer alike. Here, we demonstrate that ubiquitin-specific protease 7 (USP7)^21-32^ controls leukemia growth by stabilizing the levels of the NOTCH1 and JMJD3 demethylase. USP7 is overexpressed T-ALL and is transcriptionally regulated by NOTCH1. In turn, USP7 controls NOTCH1 through deubiquitination. USP7 is bound to oncogenic targets and controls gene expression through H2B ubiquitination and H3K27me3 changes via stabilization of NOTCH1 and JMJD3. We also show that USP7 and NOTCH1 bind T-ALL superenhancers, and USP7 inhibition alters associated gene activity. These results provide a new model for deubiquitinase activity through recruitment to oncogenic chromatin loci and regulation of both oncogenic transcription factors and chromatin marks to promote leukemia. USP7 inhibition^33^ significantly blocked T-ALL cell growth *in vitro* and *in vivo.* Our studies also show that USP7 is upregulated in the aggressive high-risk cases of T-ALL and suggest that USP7 expression might be a prognostic marker in ALL and its inhibition could be a therapeutic tool against aggressive leukemia.

## Results

Loss of the repressive mark H3K27me3 is important for leukemia initiation and progression^14^, and NOTCH1 binding to DNA and H3K27me3 loss correlate during leukemogenesis^14^. Previously, we identified and characterized the pro-oncogenic role of the epigenetic modulator Jumonji D3 (JMJD3)^34-37^, which mediates NOTCH1 oncogenic function in T-ALL^15^. JMJD3 is a NOTCH1 interactor and is recruited by NOTCH1 to oncogenic targets. We demonstrated that inhibition of JMJD3, using the small molecule inhibitor GSKJ4, yielded leukemic cytotoxicity, thereby rendering this demethylase an attractive target for pharmacologic development^15^.

To identify potential NOTCH1/JMJD3-associated proteins and understand NOTCH1 complex stability in leukemia, we retrovirally expressed hemaglutinin (HA) tagged-JMJD3 in CCRF-CEM T-ALL cells. We performed liquid chromatography-tandem mass spectrometry (LC-MS/MS) following immunoprecipitation (IP) of HA-JMJD3 (Fig. 1a). 101 proteins were uniquely associated with HA-JMJD3, and these proteins were enriched in members of the ubiquitin specific protease (USP) family of deubiquitinases (DUBs), including USP7, USP9X, USP24, and USP47 (Fig. 1b, Supplementary Table 1, and Supplementary Fig. 1a). Indeed, gene ontology analysis of the JMJD3 interactome showed enrichment in protein metabolic processes and members of the proteasome complex (Supplementary Fig. 1b).

**Figure 1.**
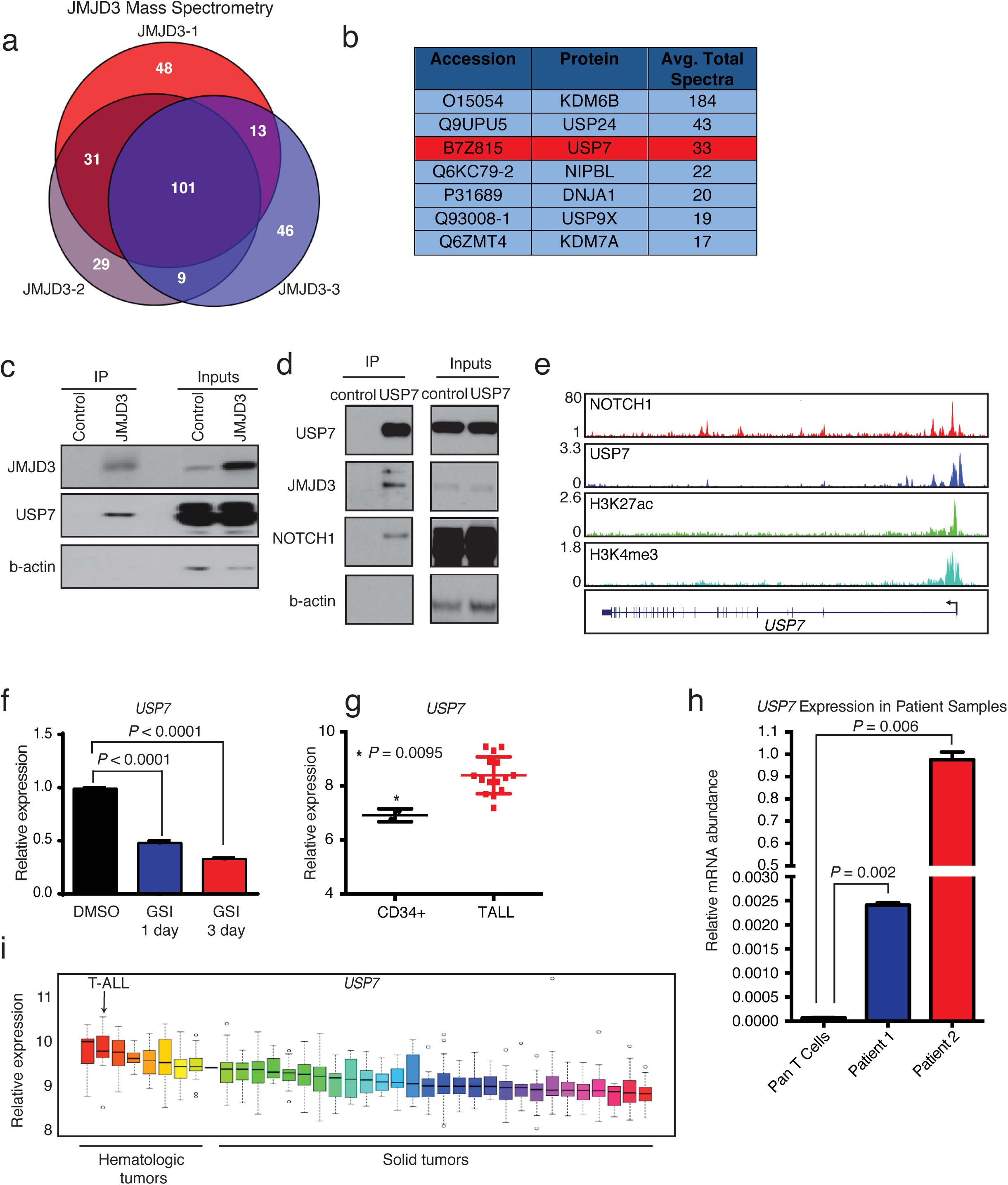
USP7 is a member of the JMJD3/NOTCH1 complex in T-ALL. **a,** HA immunoprécipitation (for HA-tagged JMJD3) followed by mass spectrometry in CCRF-CEM T-ALL cells. Shown is the overlap of HA-JMJD3-asssociated proteins across 3 biological replicates, revealing 101 common proteins associated with HA-JMJD3. **b,** Top interactors for JMJD3, showing the average spectral count of 3 technical replicates. Members of the USP family are among these top interacting proteins. **c,** Representative Western blots following immunoprecipitation of HA-JMJD3 in CCRF-CEM T-ALL cells. Control cells express the HA-tagged vector. **d,** Western blots following immunoprecipitation of USP7 in CUTLL1 T-ALL cells, showing interactions of JMJD3 and NOTCH1 with USP7. **e,** ChIP-Seq tracks showing NOTCH1 binding in CUTLL1, and USP7, H3K27Ac, and H3K4me3 in JURKAT T-ALL cells, at the *USP7* gene. **f,** Relative mRNA levels of *USP7* in CUTLL1 T-ALL cells following treatment with DMSO or 1 μM GSI for 1 and 3 days. Error bars represent the mean ± SD of 3 technical replicates. **g,** Relative mRNA expression of *USP7* in normal human CD34^+^ (*n*=2) and patient T-ALL bone marrow cells (*n*=15) (GSE62144), showing the mean ± SD. **h,** Relative *USP7* mRNA expression in normal T cells (pan T cells) and primary human T-ALL cells. Error bars represent the mean ± SD of technical duplicates. **i,** Relative mRNA expression of *USP7* in different tumor types from the Cancer Cell Line Encyclopedia. RMA = Robust Multi-array Average.

Dynamic NOTCH1 (de)ubiquitination plays a pivotal role in the regulation of NOTCH1 protein levels in leukemia, and ubiquitin ligases such as FBXW7 have been demonstrated to regulate NOTCH1 degradation and are tumor suppressors in T-ALL^38,39^. To this end, we hypothesized that USPs might be acting as oncogenic cofactors for the NOTCH1 complex, controlling complex stability and, ultimately, transcriptional potency. Ubiquitination occurs through a sequence of enzymatic reactions^40^ involving the activity of two E1 ubiquitin-activating enzymes, multiple E2 enzymes and hundreds (~700) of E3 enzymes in humans. E3 ligases determine complex specificity. In contrast, DUBs^41,42^ mainly include the ubiquitin-specific protease superfamily (USP/UBP, 58 members), the ovarian tumor (OUT, 14) superfamily, the Machado-Josephin domain (MJD, 5) superfamily, the ubiquitin C-terminal hydrolase (UCH, 4) superfamily (all the aforementioned categories are cysteine proteases) and the Jab1/Mov34/Mpr1 Pad1 N-terminal+ (MPN+) (JAMM, 14) domain superfamily proteins that binds zinc (metalloproteases).

USP7^21-32^ was the second most abundant interacting partner of JMJD3. We confirmed interaction between NOTCH1, JMJD3 and USP7 in reciprocal IP experiments (Fig. 1c,d)^43^. NOTCH1 directly binds the *USP7* gene locus (Fig. 1e) and inhibition of the NOTCH1 pathway, using gamma secretase inhibitor (GSI), led to downregulation of *USP7* and the well-characterized NOTCH1 target *HES1* (Fig. 1f and Supplementary Fig. 1c). RNA expression analysis showed that USP7 was significantly overexpressed in T-ALL compared to normal CD34+ cells and T cells, as well as other hematologic and solid tumors, correlating with expression of *HES1* (Fig. 1g-i and Supplementary Fig. 1d).

We hypothesized that USP7 might control NOTCH1 protein levels, as USP7 and NOTCH1 protein levels showed a positive correlation in a panel of T-ALL cell lines and leukemia samples (Fig. 2a-c). Overall, USP7 was substantially expressed in all T-ALL cell lines tested (Supplementary Fig. 1e). We genetically silenced USP7 in T-ALL using CRISPR-Cas9 genome editing (Supplementary Fig. 2a) and analyzed the growth of T-ALL cells. Upon deletion of USP7 using two different sgRNAs, there was significant inhibition of T-ALL cell growth (Fig. 2d). We also examined NOTCH1 protein levels upon expression of sgRNAs against USP7, and observed a decrease in the levels of NOTCH1, suggesting that USP7 might control NOTCH1 protein stability (Supplementary Fig. 2b). USP7 silencing led to increased apoptosis of T-ALL cells, as demonstrated using shRNA against USP7 (Supplementary Fig. 2c,d).

**Figure 2.**
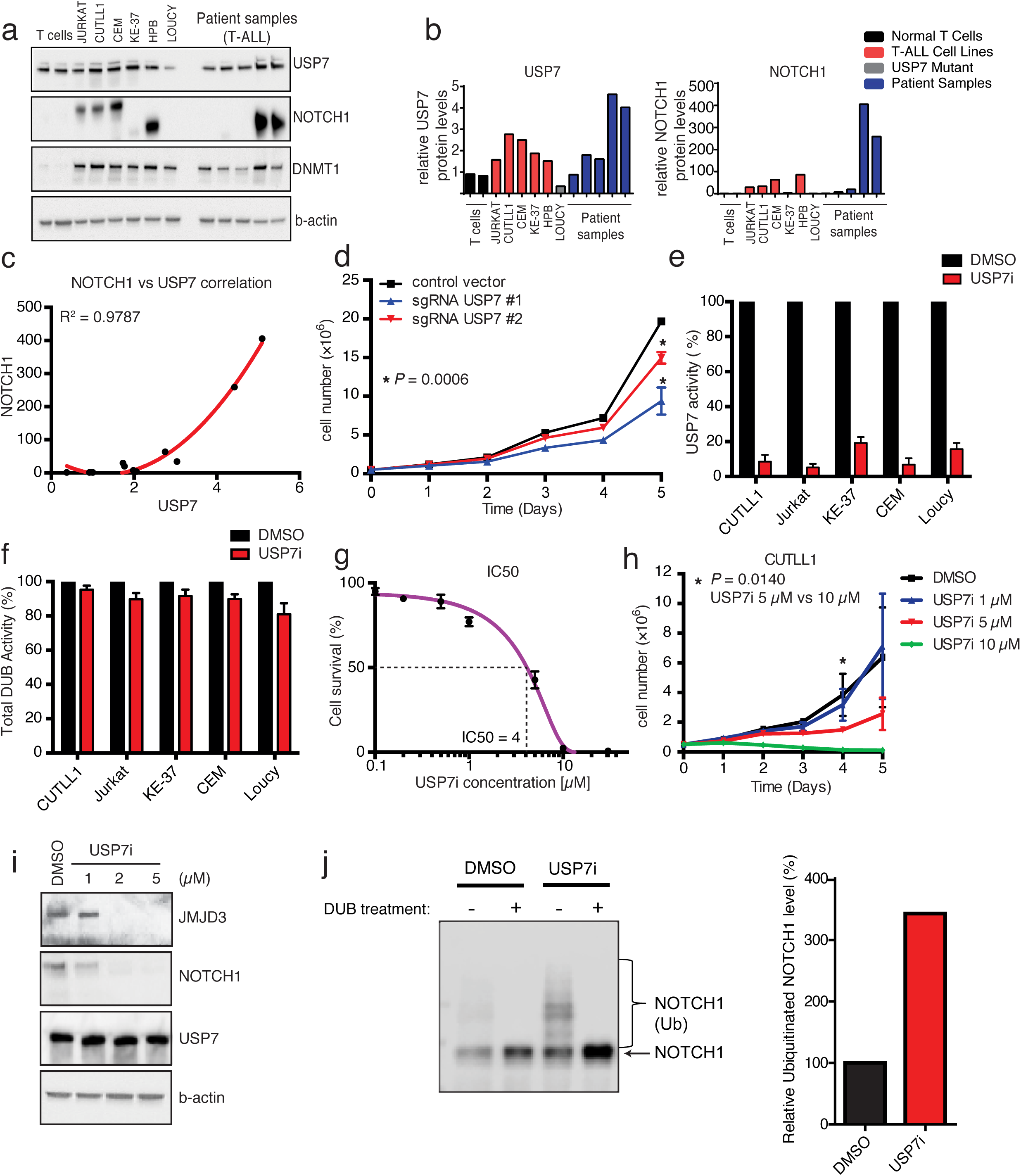
USP7 is a novel deubiquitinase for N1-IC in T-ALL. **a,** Western blots show levels of USP7, NOTCH1, and DNMT1 (positive control) in a panel of T-ALL cell lines (*n*=6) and primary patient samples (*n*=5) compared to different normal primary T cell samples (*n*=2). **b,** Quantification of panel a. **c,** A second order polynomial curve was fitted to the data in panel **b,** where R^2^=0.9787. The Spearman’s rank correlation test was applied to determine the correlation between NOTCH1 and USP7 (*P*=0.0009218). **d,** JURKAT T-ALL cell growth, as determined by trypan blue staining, in cells expressing doxycycline-inducible Cas9 and either a control vector or one containing a USP7-specific single guide RNA (sgRNA). Cells were treated with 1 μg /ml doxycycline for 3 weeks prior to the start of the experiment. Shown is the mean ± SD of technical triplicates, representative of 2 independent experiments. **e,** Treatment of a panel of human T-ALL lines with P217564 inhibitor (USP7i) for USP7 catalytic activity. Results have been normalized based on the DMSO control study. **f,** Shown is the effect of USP7i treatment on total DUB activity in the same samples shown in panel e. g, IC50 curve of USP7i in CUTLL1 T-ALL cells. **h,** Growth curve of CUTLL1 T-ALL cells upon daily treatment with increasing concentrations of USP7i. Shown is the average of technical duplicates from a combination of 2 individual experiments. **i,** Western blot analysis of JMJD3, NOTCH1, and USP7 protein levels upon treatment of JURKAT T-ALL cells with increasing concentrations of USP7i for 24h. Representative of 4 individual experiments. **j,** Western blot showing NOTCH1 ubiquitination, as determined by the UbiTest assay (LifeSensors), upon treatment of JURKAT T-ALL cells with DMSO (control) or USP7i for 8h. DUB treatment resulted in removal of ubiquitin. Quantification of ubiquitinated NOTCH1 is shown on the right.

Based on our findings that USP7 might regulate NOTCH1 oncogenic complexes, we sought to exploit USP7 inhibition as a therapeutic tool in leukemia. We took advantage of a new generation of USP7-specific inhibitors (USP7i), developed by Progenra, successfully used in preclinical models of multiple myeloma and neuroblastoma and in immunological contexts^33,44,45^. USP7i dramatically inhibited USP7 activity (up to 90% inhibition) without significantly blocking total DUB activity in T-ALL (Fig. 2e,f and Supplementary Fig. 3a-c). All T-ALL lines tested had substantial USP7 protein levels and various but significant enzymatic activities (Supplementary Figs. 1e, 3d and Fig. 2a, b). These compounds significantly inhibited leukemic cell growth in the μM range of concentrations (Fig. 2g,h and Supplementary Fig. 4a, IC50 = 4 μM).

To further delineate the mechanism behind this growth inhibition, we treated cells with USP7i for one and 3 days and performed apoptosis and cell cycle analysis, using Annexin V and DAPI staining, respectively. Although there was a significant increase in apoptosis upon treatment with USP7i (Supplementary Fig. 4b), there was no significant effect on cell cycle (Supplementary Fig. 4c). These results were comparable to effects on cell growth upon silencing of USP7 (Supplementary Fig. 2d).

We next measured NOTCH1 protein levels upon treatment with USP7i to evaluate the role of USP7 on NOTCH1 stability. We observed a significant decrease in the levels of NOTCH1 and JMJD3 upon inhibition of USP7 (Fig. 2i) starting at 1uM USP7i. To confirm the role of USP7 in NOTCH1 stabilization, we expressed wild-type USP7 together with NOTCH1, and measured NOTCH1 expression in 293T cells. Indeed, NOTCH1 was stabilized in the presence of USP7 (Supplementary Fig. 4d). To evaluate the role of USP7 in NOTCH1 ubiquitination, we inhibited USP7 catalytic activity in JURKAT T-ALL cells, pulled down ubiquitinated proteins using tandem ubiquitin binding entity (TUBE) technology^46^, and measured NOTCH1 levels (Fig. 2j). Indeed, NOTCH1 ubiquitination was increased upon USP7 inhibition. JMJD3 overexpression using a retroviral system could not rescue the effect of the drug on leukemic cells (Supplementary Fig. 4e), potentially due to the simultaneous requirement for high levels of NOTCH1 expression for leukemia maintenance.

Due to the activity of intracellular NOTCH1 (N1-IC) on chromatin, we evaluated genome-wide USP7 binding via chromatin immunoprecipitation combined with sequencing (ChIP-Seq). We identified ~10,000 high-confidence peaks for USP7 that localized near promoters (Fig. 3a) and had a strong overlap with NOTCH1 binding sites (Fig. 3b). Both USP7 and NOTCH1 bound to classical NOTCH1 targets, such as *DTX1* and *NOTCH3,* as well as *NOTCH1* itself (Supplementary Fig. 5a). Pathway analysis of the ChIP-Seq data showed a striking enrichment for NOTCH1 signaling (Fig. 3c). Furthermore, NOTCH1 recruited USP7 to chromatin, as treatment with GSI significantly depleted chromatin-bound USP7 levels (Fig. 3d). Global cellular levels of USP7 did not change over the period of treatment (Fig. 3d), due to the stability of USP7, as we showed in cycloheximide (protein synthesis inhibitor) studies (Supplementary Fig. 5b). These findings suggest that USP7 and NOTCH1 form a positive feedback loop, where NOTCH1 induces *USP7* gene expression and then recruits USP7 to chromatin, resulting in increased expression of NOTCH1 targets, through stabilization of the NOTCH1 complex.

**Figure 3.**
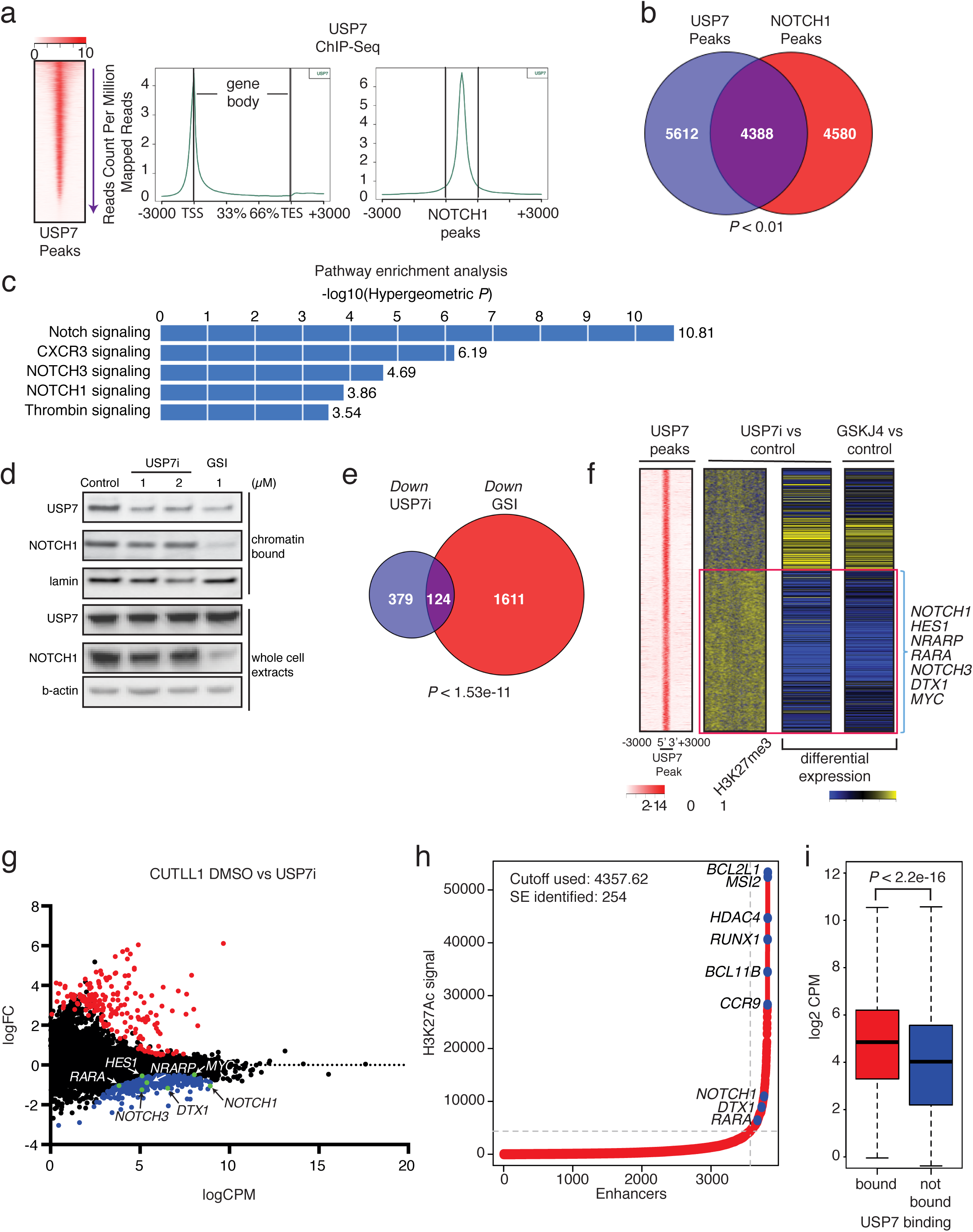
USP7 regulates NOTCH1 oncogenic targets and binds to superenhancers. **a,** Heatmap of USP7 peak intensity (left) and metaplots showing the average counts per million of USP7 ChIP-Seq signal across all genes (middle), and NOTCH1 peaks (right). **b,** Analysis of ChIP-Seq data showing the overlap of USP7 peaks with NOTCH1 peaks. **c,** GREAT (Genomic Regions Enrichment of Annotations Tool) analysis showing enrichment in gene function of the USP7-bound peaks in panel B. **d,** Representative protein levels of USP7 and NOTCH1 in the chromatin-bound fraction and whole cell extracts of JURKAT T-ALL cells upon treatment with DMSO (control), USP7i, or GSI. **e,** Analysis of RNA-Seq data showing the overlap of downregulated genes upon USP7 inhibition and GSI treatment in CUTLL1 T-ALL cells. **f,** Heatmap representation of the 10,0 strongest USP7-bound peaks, the log2 fold change of H3K27me3 upon USP7i versus DMSO treatment, and the associated log2 fold changes in gene expression upon USP7 inhibition (USP7i) and JMJD3 inhibition (GSKJ4) are shown (red box delineates cluster of downregulated genes, yellow denotes an increase in expression and H3K27me3, whereas blue indicates a decrease). NOTCH1 target genes are enriched in the downregulated cluster. **g,** Gene expression data from CUTLL1 RNA-Seq showing up‐ and downregulated genes (red and blue dots, respectively), as well as genes with no statistically significant (*P*<0.05) expression changes. NOTCH1 target genes are indicated in the plot. **h,** Superenhancer analysis using H3K27Ac ChIP-Seq data from JURKAT T-ALL cells. The y-axis shows H3K27Ac signal at the superenhancer locus and the x-axis shows the superenhancers ranked from lowest signal to highest, dots marked blue correspond to the top five SEs and NOTCH1 target SEs. **i,** Boxplot showing the difference in gene expression measured as log2 CPM (counts per million) of genes bound by USP7 (red) and not bound by USP7 (blue). Genes were considered to be bound or not bound by determining the nearest gene to a USP7 peak. There was a statistically significant increase in gene expression among bound genes versus those not bound by USP7 (*P*<2.2e-16, *t*-test).

Our data suggest that USP7 regulates important NOTCH1-driven oncogenic complexes. To focus on NOTCH1-related molecular responses we used 1 μM USP7i, as NOTCH1 levels are sensitive to this inhibitor concentration (Fig. 2i). To delineate the molecular effects of USP7 activity, we performed global expression analysis using RNA-Seq upon treatment with USP7i, GSI (NOTCH1 inhibitor), and GSKJ4 (JMJD3 inhibitor). The transcriptional signatures of NOTCH1, USP7, and JMJD3 chemical inhibition overlapped significantly, both with regards to downregulated as well as upregulated genes (Fig. 3e and Supplementary Fig. 6). Specifically, inhibition of USP7 led to downregulation of NOTCH1 targets, including *NOTCH3* and *DTX1* (Fig. 3f, g and Supplementary Fig. 7a). Similar to enzymatic inhibition, *USP7* silencing led to downregulation of NOTCH1 targets (Supplementary Fig. 7b). Gene set enrichment analysis upon USP7 inhibition showed that major oncogenic pathways (NOTCH1, MYC, DNA damage response, and metabolism) were downregulated (Supplementary Fig. 8). Together, these data show that USP7 enzymatic activity is important for sustaining oncogenic activity in T-ALL. It is well known that NOTCH1 regulates oncogenic enhancers^47,48^. Analysis of USP7 ChIP-Seq data showed that leukemia-specific superenhancers (SEs)^49,50^, as determined by H3K27 acetylation signal intensity (Fig. 3h), were enriched for binding of USP7 and NOTCH1: 192 out of 254 leukemia SEs were co-bound by USP7 and NOTCH1. In agreement with the association of SEs with transcriptional activation, we found that the USP7-bound genes were more highly expressed compared to the rest of the transcriptome (Fig. 3i) and present with lower expression upon USP7i treatment. Analysis of ChIP-Seq data for T-ALL-related transcription factors showed that USP7 binding overlapped with the binding of several of those factors (Supplementary Fig. 9). Motif analysis of USP7 binding sites showed binding motifs for some of these same factors (Supplementary Fig. 10a). Indeed, our studies using combinatorial inhibition of the bromodomain proteins that recognize acetylated histones (JQ1)^51,52^ and USP7 showed a synergistic effect (Supplementary Fig. 10b). This was further underlined by our targeted and genome wide analysis of USP7 interactions showing that USP7 interacts with Mediator 1, a main component of SEs (Arcipowski and Ntziachristos, unpublished), suggesting that USP7 may regulate gene activation via interactions with NOTCH1 and potentially other members of the T-ALL transcriptional machinery in SE areas.

NOTCH1 activity leads to eviction of the repressive mark H3K27me3 from the promoters of oncogenic targets^14,15^. We hypothesized that inhibition of USP7 would lead to an increase in H3K27me3 levels through reduction of JMJD3 protein levels. We assessed levels of H3K27me3 on USP7 targets upon USP7i treatment for 24h. Indeed, USP7 target loci that were downregulated upon USP7i treatment presented a significant gain of H3K27me3 (Fig. 3f). As USP7 has been shown to control histone ubiquitination^53,54^, we also examined histone H2B lysine 120 ubiquitination (H2Bub) levels following USP7 inhibition. We showed that USP7 colocalized with genes exhibiting a gain of H2B ubiquitination upon USP7 inhibition (Supplementary Fig. 11a). Interestingly, these downregulated genes exhibited a striking loss of H3K79me2 and Pol II occupancy (Supplementary Fig. 11b) in agreement with what was also shown previously in different mammalian contexts, such as in stem cell biology^55,56^. These changes in HBub and H3K27me3 seemed to be localized to USP7 targets and not a genome-wide phenomenon as global levels of these marks were not significantly altered following treatment with USP7i as we demonstrated using global bottom-up proteomics^57^ (Supplementary Fig. 11c). We also noticed a focal gain of H2A lysine 119 ubiquitination (H2Aub) on those down-regulated loci, potentially associated to H3K27me3 changes (Supplementary Fig. 11a)^58^. To better understand whether gene expression depends on both H3K27me3 demethylation and H2B deubiquitination changes and whether levels of those two marks depend on each other, we tried to uncouple those two processes by treatment of JURKAT TALL cells with USP7i, followed by wash-off the inhibitor and treatment with DMSO (control) or GSKJ4 JMJD3 inhibitor. In principle, removal of USP7i should lead to a fast and significant increase in gene expression in the case of DMSO (Supplementary Fig. 11d). In contrast to DMSO, we should see only a partial derepression of gene expression upon GSKJ4 treatment, if USP7 affects H2Bub independently of changes in H3K27me3. Indeed, GSKJ4 treatment compromised expression of NOTCH1 targets compared to the DMSO control (Supplementary Fig. 11d) showing that USP7 controls H2Bub and H3K27me3 independently.

This significant effect of USP7i on leukemia cells led us to further explore USP7 levels in high-risk TALL cases and compare it to low‐ and medium-risk T-ALL cases. To answer this, we used reverse phase protein array (RPPA) analysis^59,60^ for NOTCH1 and USP7 protein levels using a panel of 64 pediatric leukemia patient samples. We noticed that USP7 and NOTCH1 levels significantly correlate in T-ALL (Fig. 4a) further reinforcing the idea of the positive feedback loop in leukemia. Using data for 50 available patient samples, we demonstrate increased levels of USP7 in high-risk disease compared to medium‐ or low-risk disease (HR (15 patients) vs non-HR (35 patients, Fig. 4b)). Risk is calculated based on minimal residual disease (MRD) at days 35 and 78 post initiation of chemotherapy. Overall MRD levels are the most reliable and recognized markers of drug resistance, thus the most important parameter to consider for therapeutic treatments. These findings show that USP7 inhibition is a valid therapeutic tool in high-risk leukemia, where other treatments have failed to lead to significant disease regression. Our findings also demonstrate a potential role for USP7 as a prognostic marker in T-ALL. This significant finding led us to assess the potential of USP7 inhibition alone or in combination with JMJD3 to inhibit leukemia growth *in vivo,* we evaluated inhibitor toxicity by injecting mice intravenously (i.v.) with vehicle, USP7i alone, or USP7i with increasing concentrations of GSKJ4. The GSKJ4 concentrations used have been previously shown to successfully inhibit tumor growth^61,62^. Immunocompromised mice treated daily for five days showed no signs of treatment-associated toxicity, as determined by complete blood cell count, body weight and analysis of spleen and liver (Supplementary Fig. 12a and data not shown). We then transplanted luciferase-expressing JURKAT T-ALL cells either intravenously (i.v., Fig. 4c) or subcutaneously (SubQ, Supplementary Fig. 12b) into immunocompromised mice^14,15^. Upon tumor detection using bioluminescence imaging (IVIS), mice were randomized into different groups that were treated with vehicle or USP7i [10 mg/kg]. In both models, tumors in USP7i-treated animals showed a significant growth disadvantage compared to the control group (Fig. 4c and Supplementary Fig. 12b). Similar results were obtained using a primagraft i.v. model, where T-ALL progression was detected in the peripheral blood using the marker human CD45 (Fig. 4d and Supplementary Fig. 12c). As H3K27me3 changes are the major epigenetic alteration in TALL progression, we and others have successfully used the JMJD3 inhibitor (GSKJ4) against T-ALL in the past^15,61^ with decent effect on leukemia inhibition. To further potentiate the therapeutic impact of USP7i in TALL, we investigated the effect of combinatorial USP7 and JMJD3 inhibition on leukemia growth by treating mice with USP7i and GSKJ4 in a subcutaneous model of T-ALL and monitored animal survival (Fig. 4e). Similarly to our previous studies (4c and Supplementary Fig. 12b, c) we found that USP7i as a single therapy significantly inhibits tumor growth (Fig. 4e, top panel) and extends mouse survival (Fig. 4e, bottom panel). Moreover, combinatorial inhibition significantly decreased tumor growth, more effectively than vehicle or USP7 inhibition alone (Fig. 4e, top/right panel) without any associated toxicity based on our analysis of mouse weight (Supplementary Fig. 12d) and histology studies (not shown). Moreover, we demonstrated significant survival differences between mice injected with vehicle or those injected with a combination of USP7 and JMJD3 inhibitors (Fig. 4e, bottom panel). Importantly, USP7i/GSKJ4 treatment yielded a prolonged mouse survival compared to vehicle or USP7i therapy (Fig. 4e bottom panel and data not shown), lending rationale to the use of combinatorial drug treatments against epigenetic regulators in cancer.

**Figure 4.**
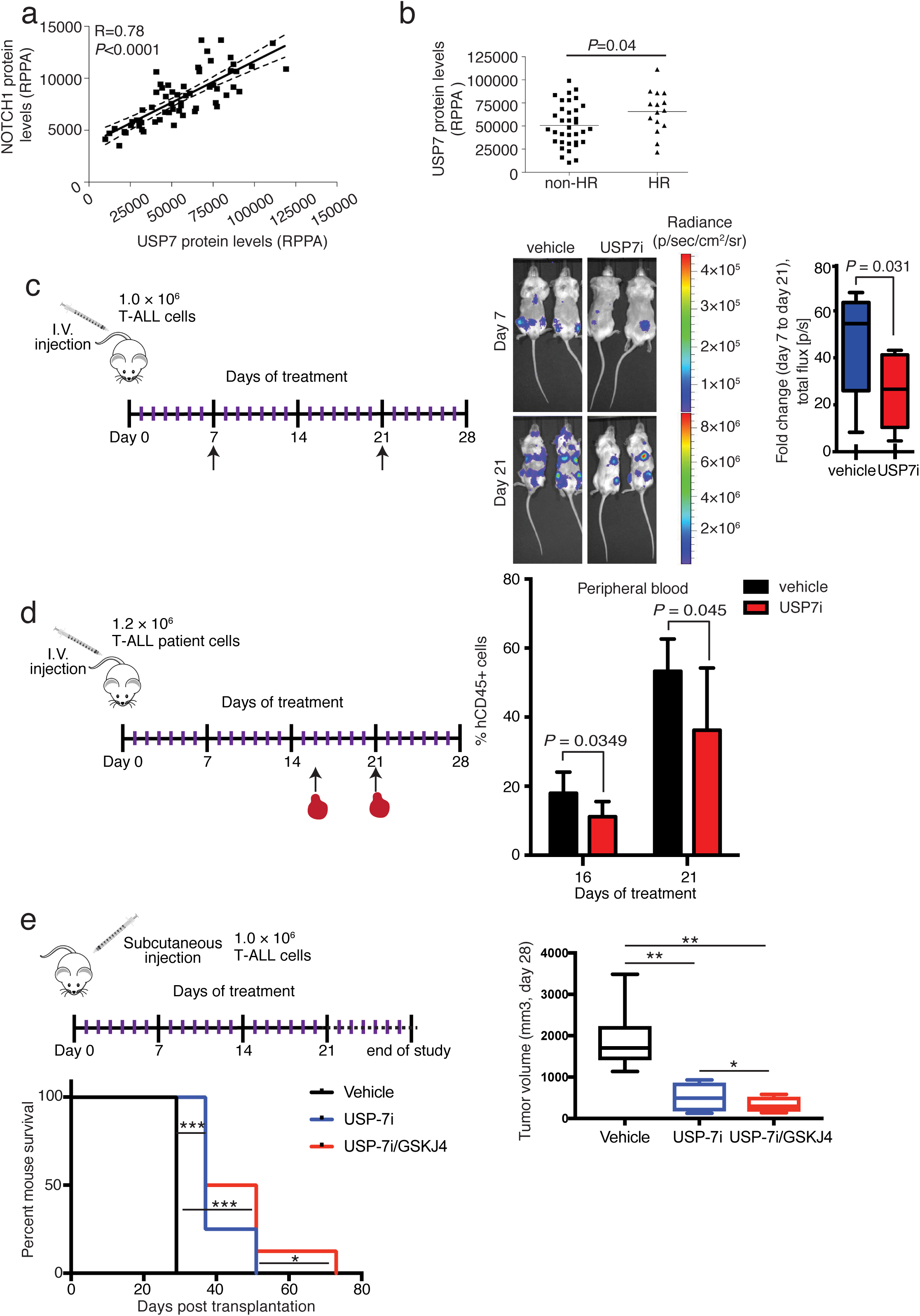
USP7 inhibition blocks leukemia growth in preclinical models of T-ALL. **a,** USP7 protein levels are correlated with NOTCH1 levels in primary T-ALL. **b,** USP7 expression is significantly associated with high-risk disease. HR: high risk. *P*<0.05 between non-HR (*n*=35) and HR (*n*=15). **c,** JURKAT T-ALL cells were injected intravenously (i.v.) into immunocompromised mice. Upon detection of leukemic blasts using bioluminescence (with IVIS), mice were treated i.p. with 10 mg/kg USP7i 3 days/week for 3 weeks. Relative luminescence intensity is shown for 2 representative mice per treatment group on days 7 and 21 of treatment. The fold change in total flux from day 7 to day 21 is shown on the right (vehicle, n=8; USP7i, n=9). **d,** Patient T-ALL cells expressing high levels of NOTCH1 and USP7 were injected i.v. into immunocompromised animals and, upon detection of human CD45+ cells in the peripheral blood, mice were administered USP7i at 10 mg/kg 3 times/week for 3 weeks. The mean percentage of hCD45+ cells in the peripheral blood ±SD at days 16 and 21 (at the end of the treatment period) is shown (*n*=7). **e,** Immunocompromised mice were injected subcutaneously with JURKAT T-ALL cells. Once tumors became visible, mice were administered USP7i i.v., alone or in combination with GSKJ4, 3 times/week till termination of the study. Shown is mouse survival (lower left panel) and the mean tumor volume ± SD (right panel) for the different groups (vehicle, *n*=7; USP7i, *n*=5; and USP7i/GSKJ4, *n*=5). * *P*<0.05, ** *P*<0.01, *** *P*<0.001.

## Discussion

Therapeutic targeting of high-risk acute lymphoblastic leukemia (ALL) has been challenging, rendering this disease an unmet clinical need. In this study, we performed a series of epigenetic, biochemical, functional and pharmacological studies to show that the deubiquitinase USP7 functionally and physically interacts with and controls NOTCH1 pathway and ultimately the oncogenic transcriptional circuitry in T cell ALL. USP7 levels can be a biomarker for high-risk ALL and inhibition of its enzymatic activity could be a valid therapeutic target in this disease.

Given the previously unrecognized importance of deubiquitination in acute leukemia progression, we characterized a novel role for USP7 in T-ALL, providing the first evidence that the ubiquitination-methylation axis can serve as an oncogenic switch and therapeutic target in T-ALL. USP7 is the first deubiquitinase identified and thoroughly characterized to regulate the NOTCH oncogenic pathway. Our studies showed that USP7, NOTCH1, and JMJD3 act in a positive feedback loop, where the NOTCH1/JMJD3 complex induces expression of USP7, resulting in USP7 recruitment to target genes (Supplementary Fig. 13). Moreover, our ongoing studies show that NOTCH1 molecules with mutant PEST domain (shown in the past to control NOTCH1 levels through interaction with FBXW7^38,39^ ligase), interact with and are deubiquitinated by USP7, showing that other parts of NOTCH1 intracellular domain control its regulation through USP7 (Jin and Ntziachristos, unpublished). USP7 recruitment leads to stabilization of the oncogenic complex and activation of targets through demethylation of H3K27 and deubiquitination of H2B, as well as changes in H3K27Ac at SEs. Intriguingly, and in contrast to the previously characterized pro-oncogenic role of USP7 in T-ALL and multiple myeloma^33^, neuroblastoma^63^, chronic lymphocytic leukemia^64^ and other solid tumors^65-67^, there are mutations affecting USP7 in pediatric cancers including TAL1-positive leukemias^68,69^. Thus, similarly to what others and we showed for another epigenetic modulator, the histone demethylase UTX, which can play dual roles in NOTCH1‐ and TAL1‐ contexts of T-ALL (oncogene or tumor suppressor)^15,61,70^, the role of USP7 might also be context-specific. These context-specific roles of the epigenetic modulators are another intriguing aspect of their biology that we can exploit to develop elegant targeted therapeutic approaches in cancer. Further research is needed to study the role of USP7 in leukemia contexts other than NOTCH1-positive T-ALL, such as TAL1-positive leukemia.

Together, our findings strongly suggest that USP proteins, and USP7, in particular, may be exploited for pharmacological inhibition in certain T-ALL patients. Combinations of targeted therapies are hailed as the future of cancer therapy, with hematological malignancies leading the way (i.e. bortezomib, lenalidomide, and dexamethasone for multiple myeloma, and rituximab and ibrutinib for chronic lymphocytic leukemia). Thus, combined inhibition of USP7 and JMJD3 may represent an attractive therapeutic approach for T-ALL, especially given that small molecules generated by Progenra are already available for preclinical studies.

## Acknowledgments

We want to thank all members of the Ntziachristos laboratory for critical review of the manuscript and their comments. This work was supported by the NIH T32CA080621-11 Oncogenesis and Developmental Biology Training Grant and Cancer Smashers Postdoctoral Fellowship (to K.M.A.), and by the National Cancer Institute (R00CA188293-02), the American Society of Hematology, the Leukemia Research Foundation, the St. Baldrick’s Foundation, and the Zell Foundation (to P.N.). This work is also supported by AIRC IG 19186 grant to G.B. High-throughput sequencing data have been deposited into Gene Expression Omnibus, accession number GSE97435. Proteomics services were performed by the Northwestern proteomics core. Developmental Therapeutics Core (DTC) is supported by Cancer Center Support Grant P30 CA060553 from the National Cancer Institute, awarded by Robert H. Lurie Comprehensive Cancer Center. We want to thank Miklos Bekes and Tony Huang (New York University) for providing USP7-expressing plasmids.

## Author Contributions

P.N. designed the study, the experiments and wrote the manuscript. K.A. designed and performed most of the experiments and wrote the manuscript. C.A.M. designed and performed the analysis of genome-wide data and wrote the manuscript. E.T.B. and C.K.C. provided bioinformatics support. Q.J., Y.Z., B.T.G.D., K.K.W., M.R.J., A.G.V., F.W., J.W., H.W., I S., P.M.T., Y.A.G., N.A.A., S.P., N.V., GB., S.G., S.A.M., E.J.R., Y.T., L.W., A.S. performed experiments and contributed ideas. P.V.V. provided materials and advice related to the study. I.K. and C.M. designed and performed xenograft luciferase experiments and inhibitor treatments and helped with ideas and concepts. J.P.B. and B.B. have performed and analyzed treatments of primary samples with USP7i. B.A., V.S., M.P., G.B. and S.B. performed and analyzed RPPA and associated gene expression studies. N.L.K. provided service and helped with the proteomics studies. B.U. performed the mass spectrometry experiment and analysis. J.W. and S.K. helped with the design and execution of USP7i treatments and manuscript preparation. J.D.C. helped with the design of experiments and writing of the manuscript. A.Sh. contributed ideas and reagents and helped with the design of the study.

## Conflict of Interest Statement

Feng Wang, Jian Wu, Hui Wang, Ivan Sokirniy, Joseph Weinstock, and Suresh Kumar are employees of Progenra, Inc., developers of the USP7i compounds used in this paper. The remaining authors have no conflicts of interest.

## Figure Legends

**Supplementary Figure. 1.**
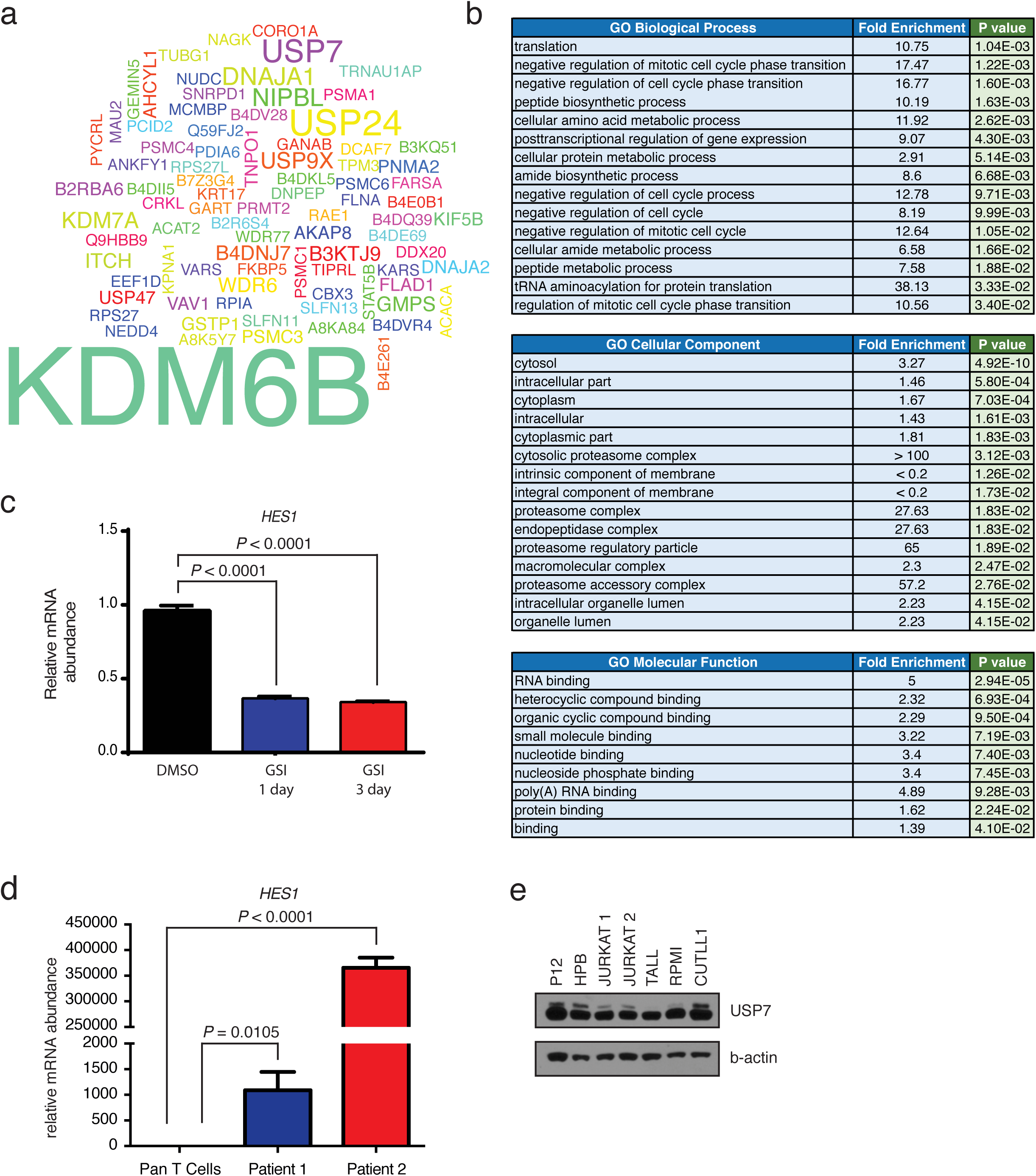
USP7 is a top interacting partner of the JMJD3/NOTCH1 complex. **a,** Word cloud of proteins (based on spectral counts) that interact with JMJD3, as determined by mass spectrometry analysis. **b,** Gene ontology analysis of the proteins in panel **a,** showing biological processes (top), cellular components (middle), and molecular functions (bottom). **c,** *HES1* mRNA levels upon GSI treatment of CUTLL1 T-ALL cells for 1 and 3 days. Error bars represent the mean ± SD of 3 technical replicates. **d,** Relative *HES1* mRNA expression in normal T cells (pan-T) and primary human T-ALL cells. Error bars represent the mean ± SD of technical duplicates. **e,** Western blot showing USP7 expression in a panel of T-ALL cell lines.

**Supplementary Figure 2.**
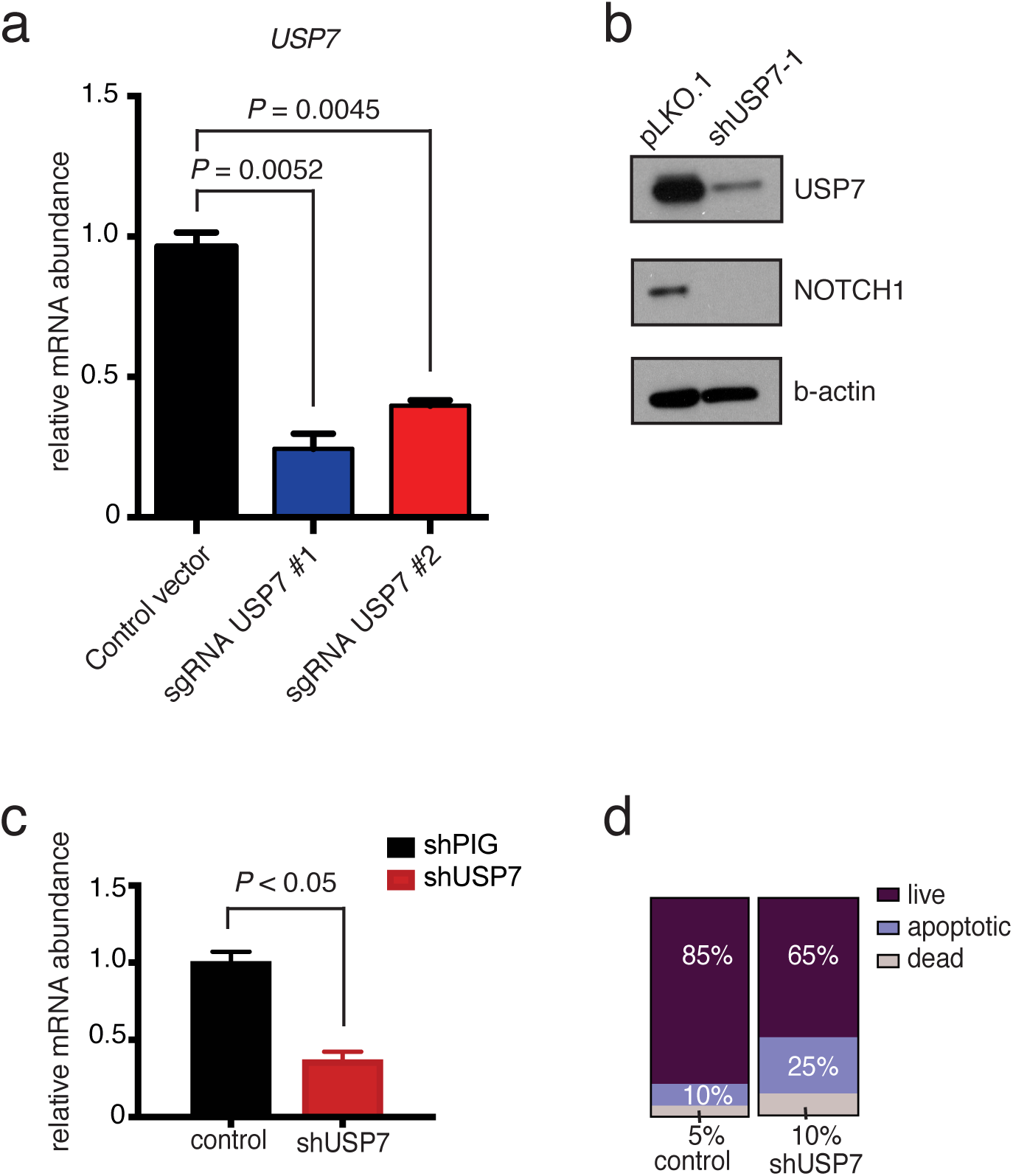
USP7 silencing leads to a reduction of NOTCH1 protein levels and increased apoptosis. **a,** Relative mRNA levels of *USP7* upon expression of USP7-specific sgRNAs. Shown is the mean ± SD of technical duplicates. **b,** Representative protein levels of USP7 and NOTCH1 upon expression of a USP7-specific sgRNA in JURKAT T-ALL cells. **c,** Knockdown of *USP7* following shRNA expression in JURKAT TALL cells. shPIG refers to the control empty vector puro-IRES-GFP. **d,** Annexin V analysis upon treatment of JURKAT cells with shUSP7 and a control retroviral vector.

**Supplementary Figure 3.**
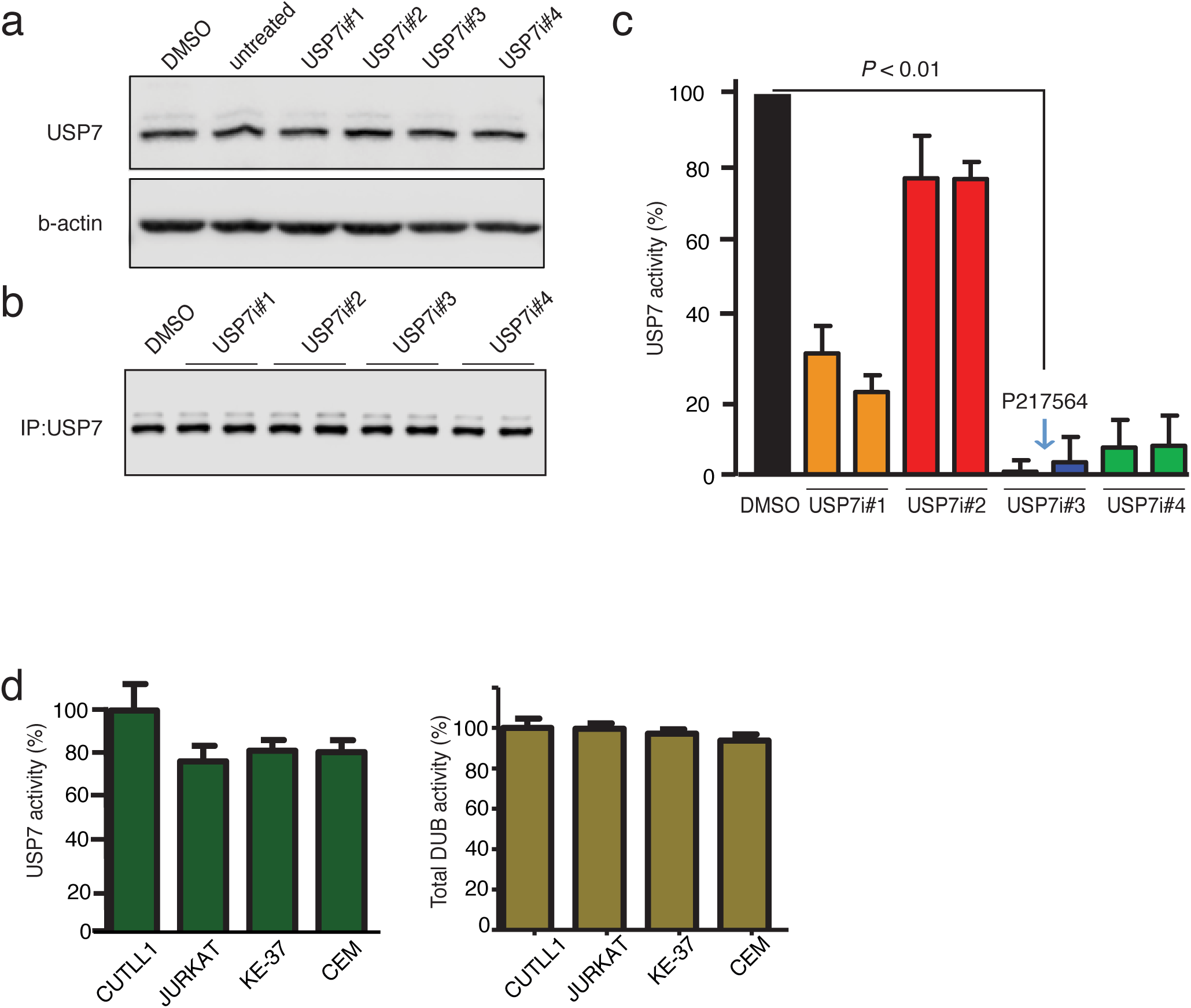
Specific inhibition of USP7 in JURKAT T-ALL. **a,** Total USP7 levels in JURKAT T-ALL cells treated with DMSO or a panel of USP7i compounds. **b,** Representative levels of USP7 upon IP of USP7 in JURKAT cells treated with DMSO or a panel of USP7i compounds. **c,** USP7 activity, as determined by ubiquitin competition, comparing the effect of DMSO (control) to a panel of 4 USP7 inhibitors. The inhibitor used in our studies, P217564, is indicated by the blue arrow. **d,** Relative USP7 and DUB activities in T-ALL cells.

**Supplementary Figure 4.**
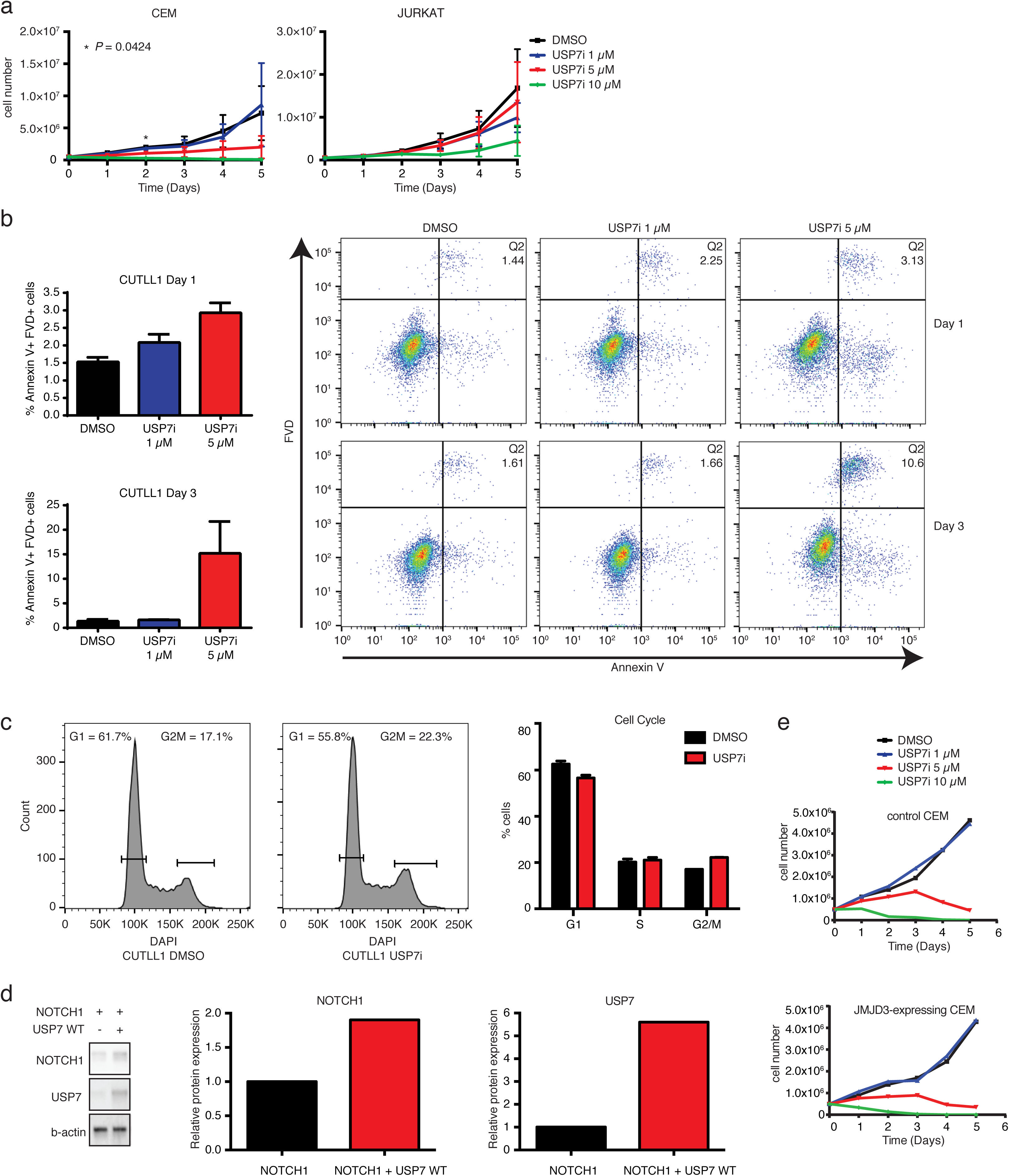
USP7 inhibition destabilizes NOTCH1 and JMJD3 and leads to leukemia cell death via apoptosis. **a,** Growth studies using CCRF-CEM (left) and JURKAT (right) T-ALL cell lines upon treatment with different concentrations of USP7i. Shown is the mean ± SD of technical duplicates, representative of 2 independent experiments. **b,** CUTLL1 T-ALL cells were treated with DMSO or increasing concentrations of USP7i for 1 or 3 days. Apoptosis was measured by Annexin V and fixed viability dye (FVD) staining. Shown is the mean ± SD of technical duplicates, and representative FACS plots showing the percentage of apoptotic cells. **c,** Shown are representative cell cycle profiles (determined by DAPI staining) of CUTLL1 T-ALL cells treated with DMSO or 1 μM USP7i for 3 days. The bar graph represents the mean ± SD of technical duplicates. **d,** A representative study showing protein levels for NOTCH1 upon expression of wild-type USP7 and NOTCH1 in 293T cells treated with the proteasome inhibitor MG132. Quantification is shown on the right. **e,** CCRF-CEM T-ALL cells were treated with DMSO or increasing concentrations of USP7i for 5 days. Shown is the growth curve of cells transfected with an empty vector (top) or HA-JMJD3 (bottom).

**Supplementary Figure 5.**
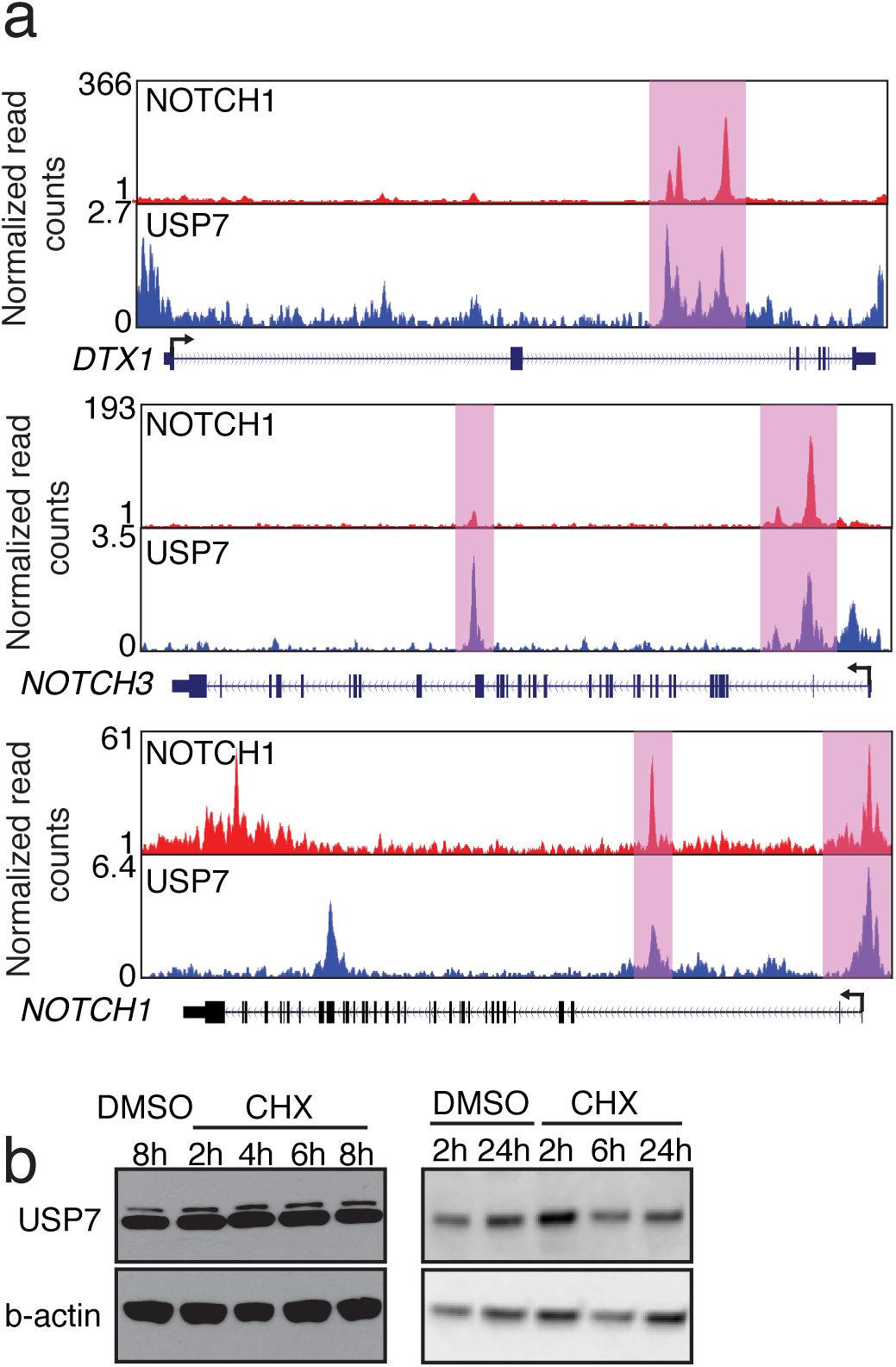
USP7 binding to NOTCH1 targets and stability of the protein. **a,** ChIP-Seq tracks showing NOTCH1 (CUTLL1) and USP7 (JURKAT) co-bound (pink highlights) at NOTCH1 targets *DTX1* and *NOTCH3,* as well as *NOTCH1.* **b,** USP7 protein levels upon treatment with 25 μg/ml cycloheximide in JURKAT T-ALL cells.

**Supplementary Figure 6.**
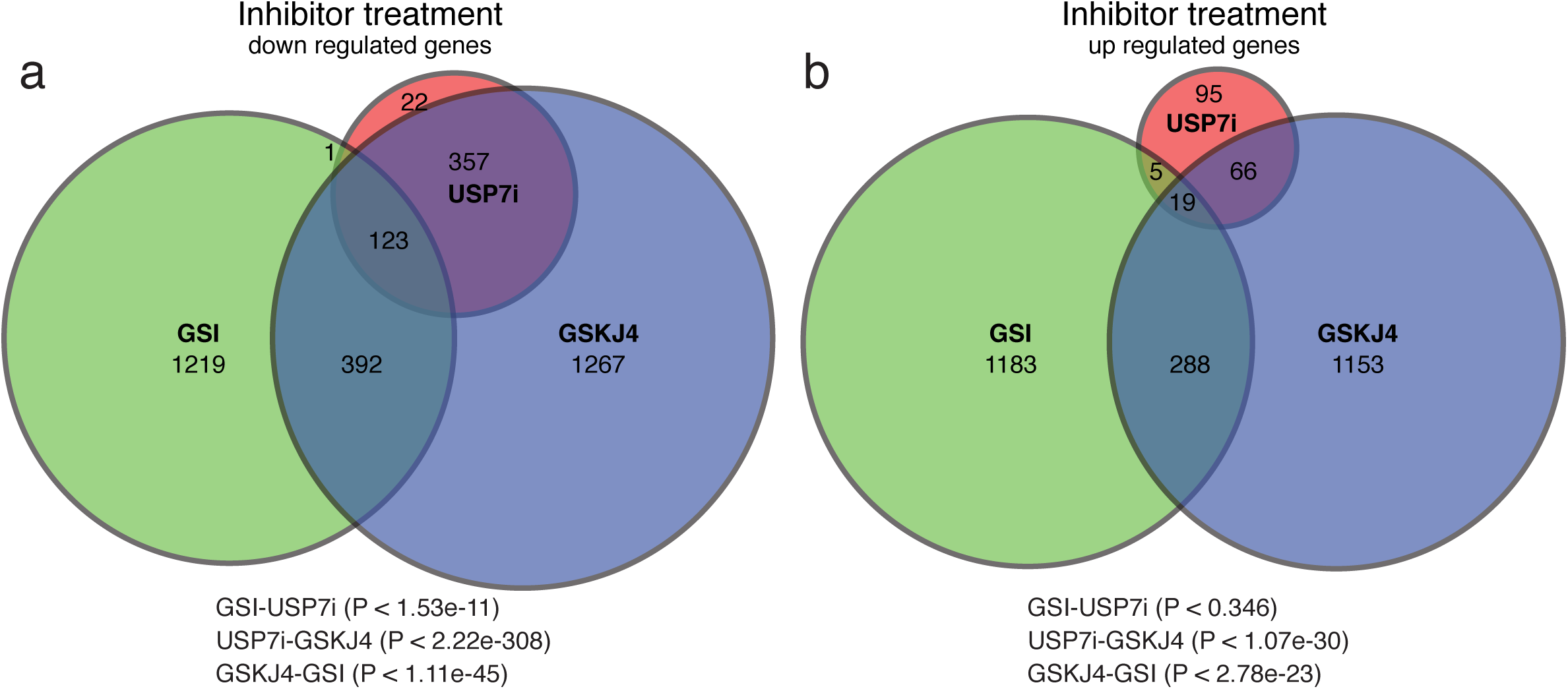
Overlap of transcriptional signatures upon USP7, NOTCH1 and JMJD3 (histone demethylase) inhibition in T-ALL cells. CUTLL1 cells were treated with USP7i, GSI, or GSKJ4 for 1 day. Shown is a Venn diagram showing RNA-seq expression of genes that are downregulated **a,** or upregulated **b,** upon USP7i, GSI, and GSKJ4 treatment.

**Supplementary Figure 7.**
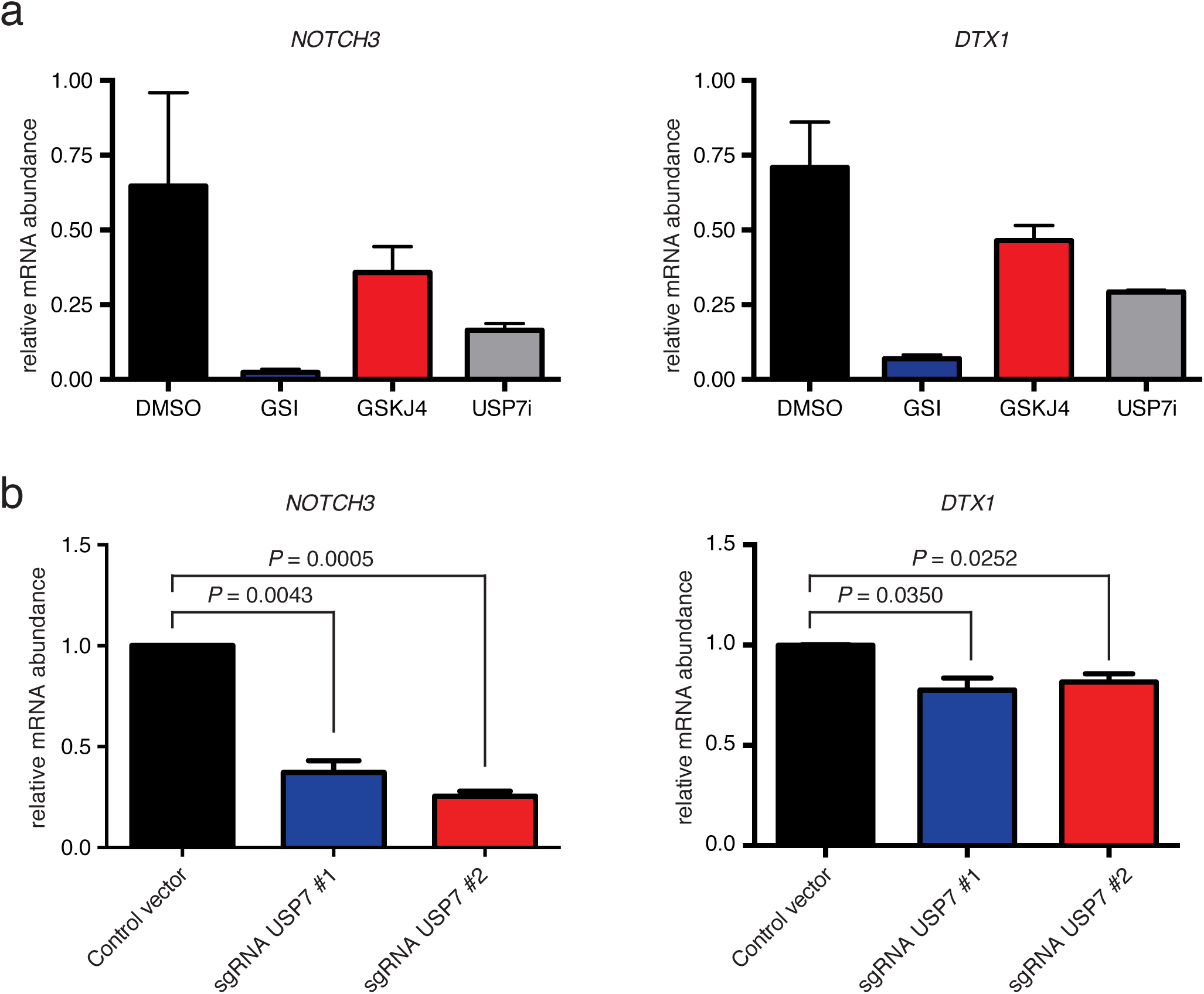
USP7 regulates NOTCH1 targets. **a,** Relative *NOTCH3* and *DTX1* mRNA abundance upon treatment of JURKAT cells with DMSO (control), or NOTCH1 (GSI), JMJD3 (GSKJ4), or USP7 inhibitors. Shown is the mean ± SD of technical triplicates. The experiment was conducted in three biological replicates. **b,** Relative mRNA levels of *NOTCH3* and *DTX1* in JURKAT T-ALL cells upon expression of sgRNAs targeting USP7. Shown is the mean ± SD of technical duplicates. The experiment was conducted in three biological replicates.

**Supplementary Figure 8.**
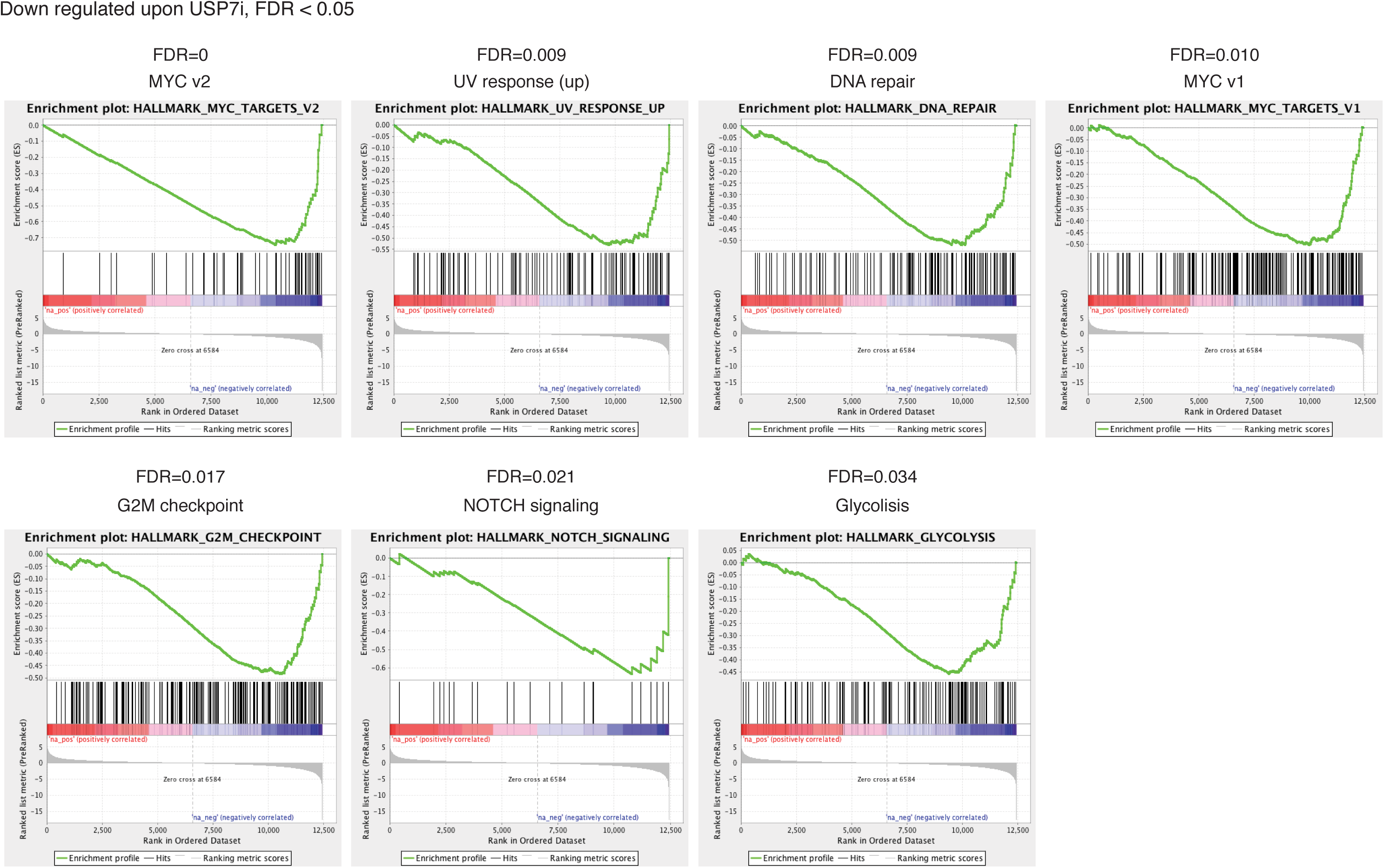
Gene set enrichment analysis for downregulated genes upon USP7 inhibition. GSEA analysis was performed on RNA-Seq data using the curated Hallmark pathway database. The False Discovery Rate (FDR) is shown for each pathway, and selected pathways with FDR<0.05 are shown. Genes in the enriched pathways were downregulated upon USP7i treatment.

**Supplementary Figure 9.**
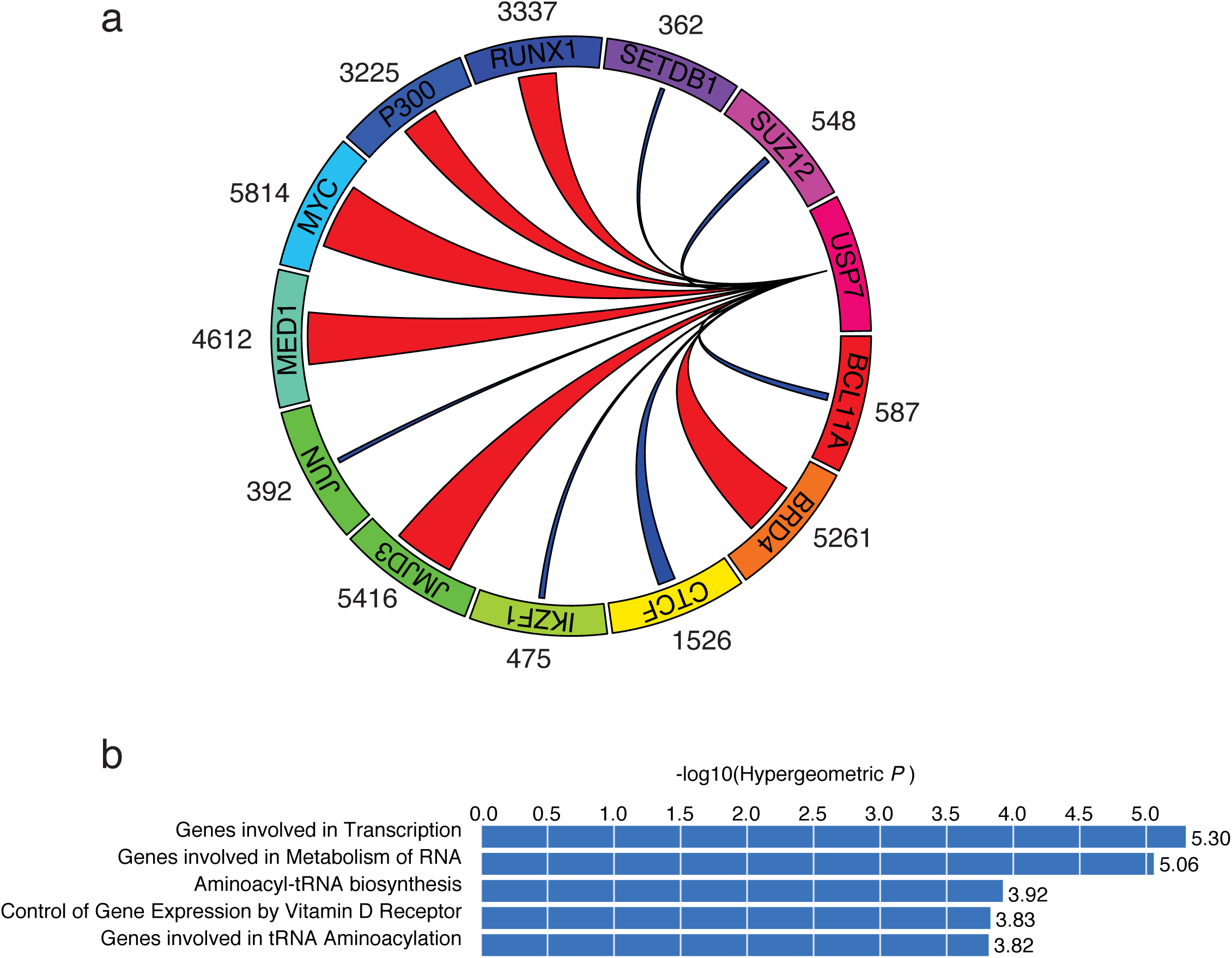
USP7 is associated with gene control elements and factors important for leukemogenesis. **a,** A circus plot is shown that represents the degree of overlap of USP7 ChIP-seq peaks with other epigenetic modulators (JMJD3, BRD4, MED1, and P300) and transcription factors (MYC and RUNX1) that play important roles in enhancer function in T-ALL. The width of the link is proportional to the number of top 10000 USP7 peaks that overlap with the top 10000 peaks of the different TFs (shown by the corresponding factor). **b,** GREAT analysis showing that regions co-bound by JMJD3, BRD4, MED1, P300, MYC, and RUNX1 are enriched in genes involved in transcriptional regulation and RNA processing.

**Supplementary Figure 10.**
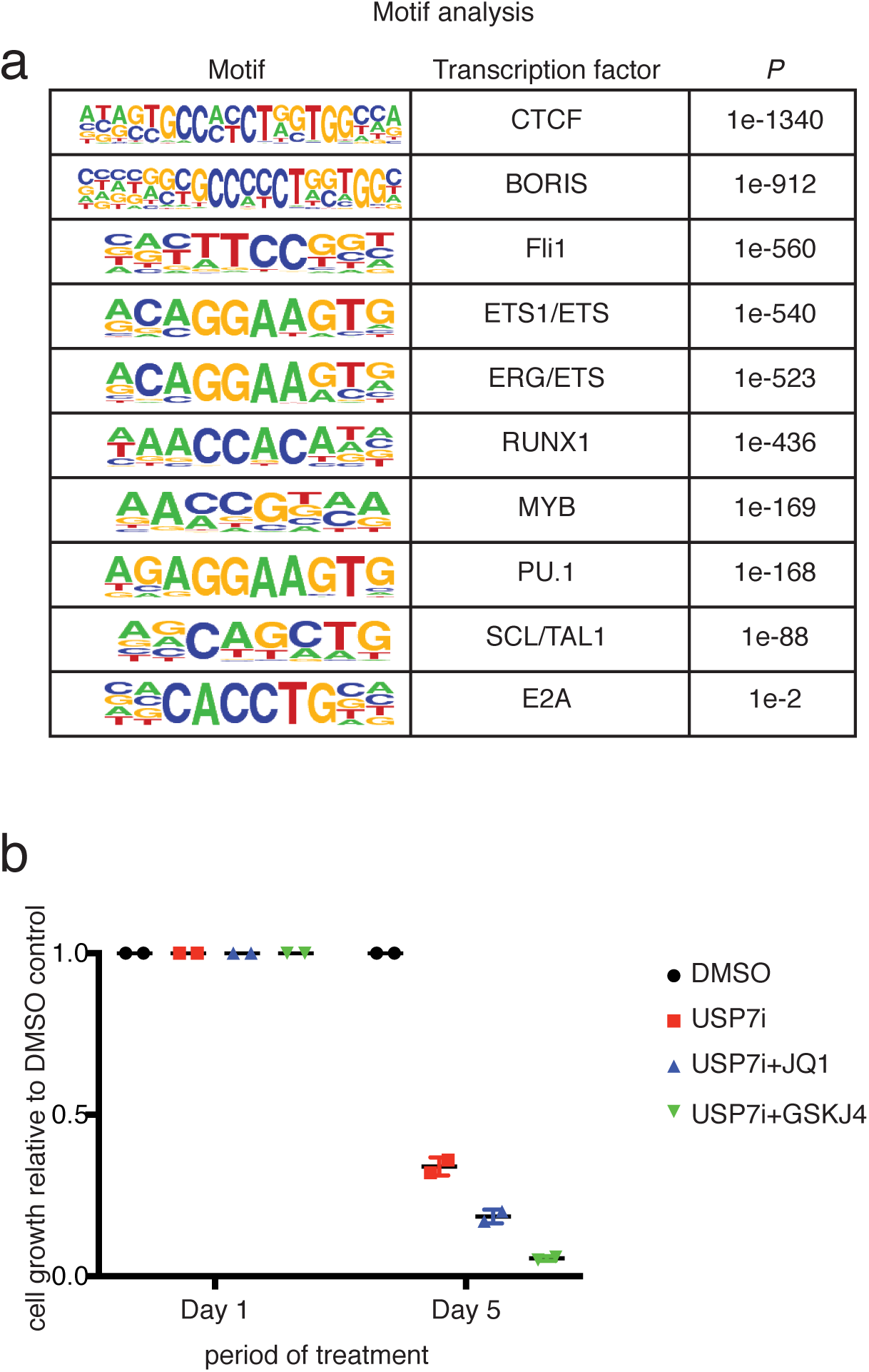
Binding sites for oncogenic transcription factors are found at sites of USP7 occupancy. **a,** Analysis of enrichment of transcription factor binding motifs in regions bound by USP7 using HOMER. **b,** Relative growth of JURKAT cells upon treatment with USP7i as a single agent or combination with JQ1 and GSKJ4 over a period of 5 days. Two technical replicates are shown.

**Supplementary Figure 11.**
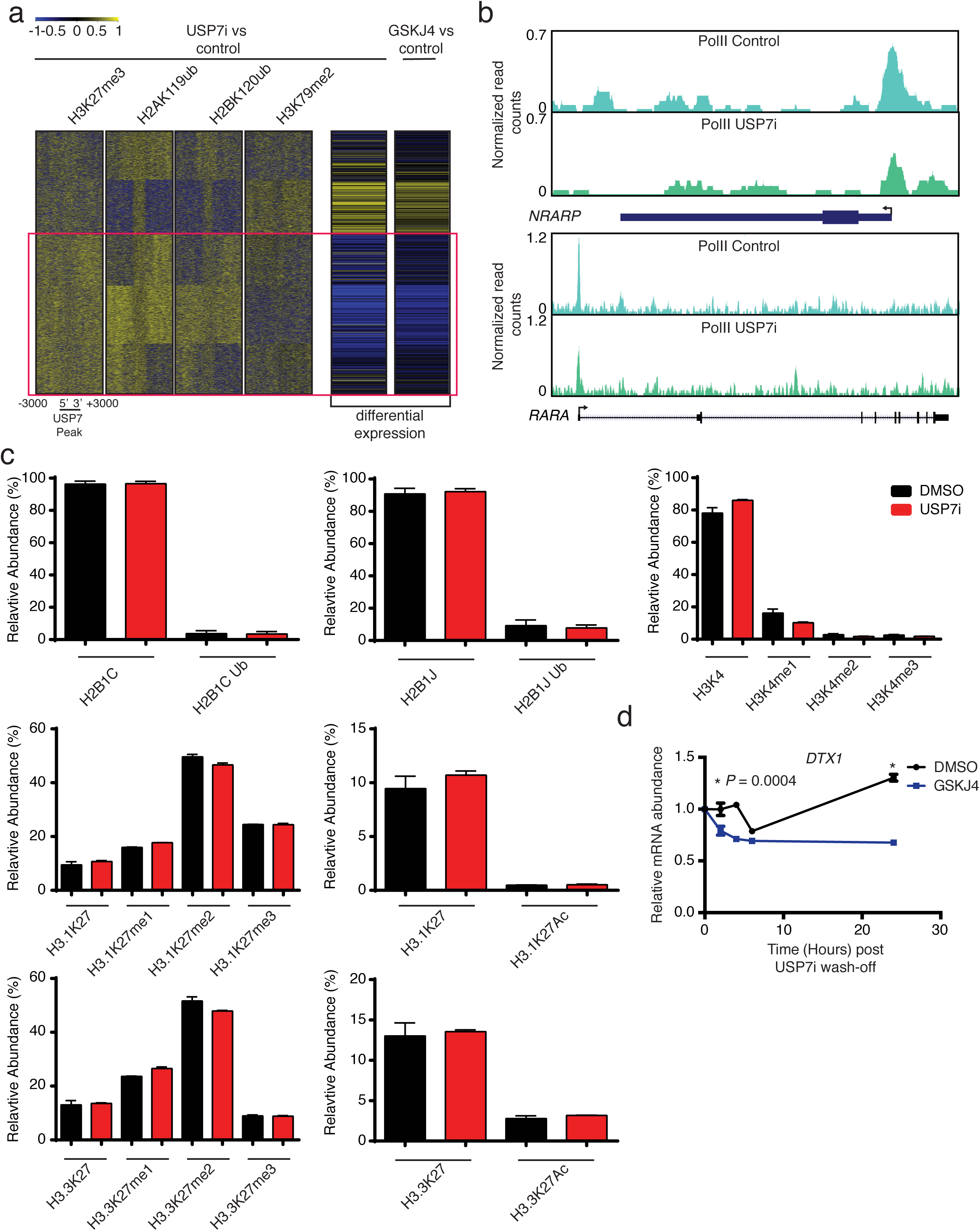
USP7 affects transcriptional regulation and Pol II binding through epigenetic changes. **a,** Heatmap showing the log2 fold change of H3K27me3, H2Aub, H2Bub, and H3K79me2 upon USP7 inhibition in JURKAT T-ALL cells, as well as the correlated log2 fold change in gene expression in CUTLL1 T-ALL cells, upon USP7 and JMJD3 inhibition (USP7i and GSKJ4, respectively). Red box delineates cluster of downregulated genes, yellow denotes an increase in expression and epigenetic marks, whereas blue indicates a decrease. Results show that H3K27me3, H2Aub, and H2Bub are increased in USP7 targets downregulated upon USP7i treatment, corresponding to lower levels of the elongation mark H3K79me2 (purple square). **b,** ChIP-Seq for Pol II in JURKAT T-ALL cells treated with DMSO or USP7i. Tracks show Pol II binding at NOTCH1 targets *NRARP* (top) and *RARA* (bottom). **c,** Bottom-up proteomics analysis for JURKAT cells upon DMSO and USP7i treatment. H3K27 methylation and acetylation, H3K4 methylation and H2B ubiquitination analysis is shown. Error bars represent ± SD of the mean of three technical replicates. **d,** JURKAT T-ALL cells were treated with USP7i for 24h, at which time the inhibitor was washed off, and cells were treated with DMSO or GSKJ4 for the indicated times. Shown are relative mRNA levels of the NOTCH1 target *DTX1.*

**Supplementary Figure 12.**
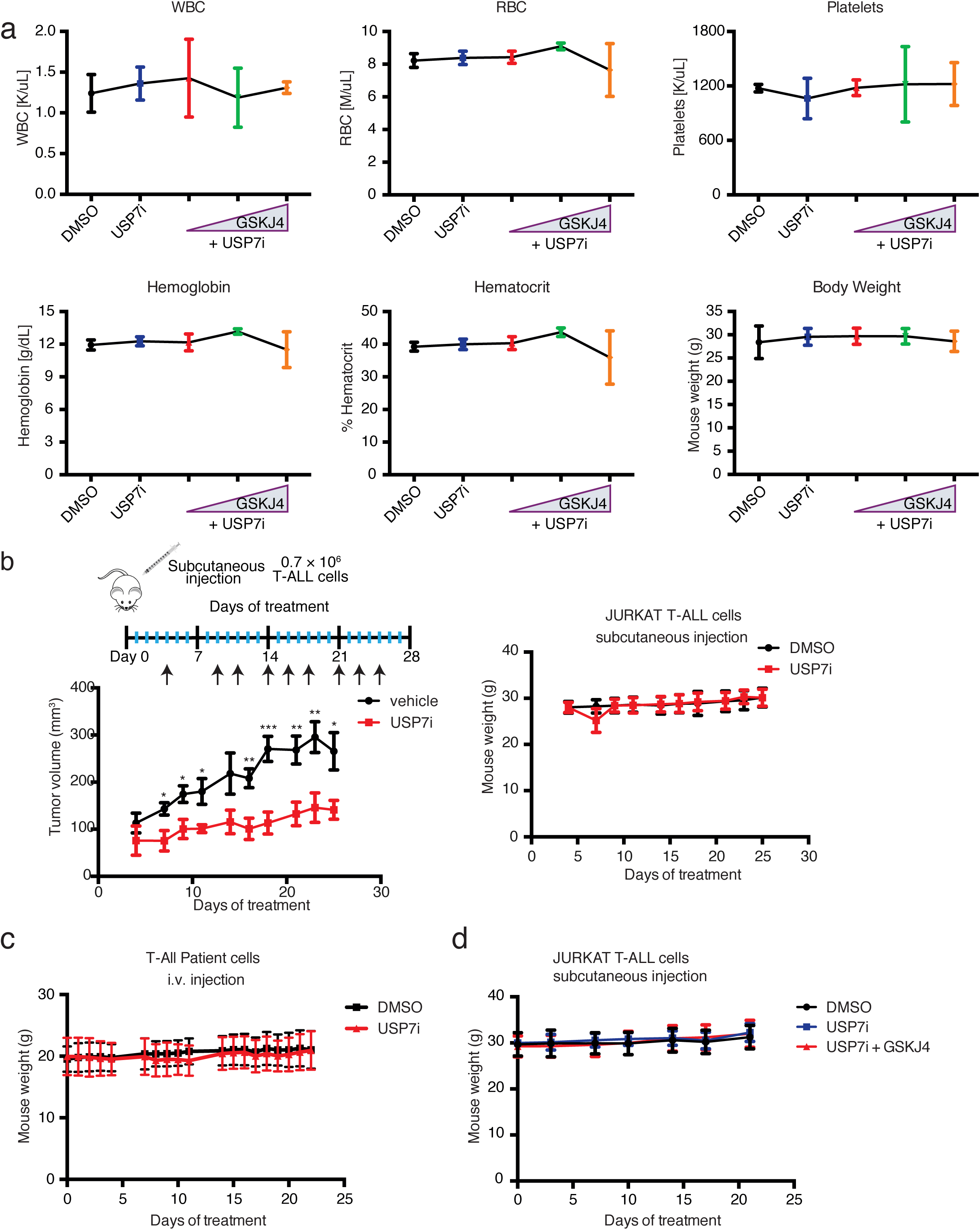
Combination treatment using JMJD3 and USP7 inhibitors is not associated with toxicity in preclinical models of T-ALL. **a,** Peripheral blood analysis and body weight upon treatment of mice with USP7i [10 mg/kg] alone or in combination with increasing concentrations of GSKJ4 at 10, 20 and 50 mg/kg (*n*=3 per treatment group) for 5 days. WBC = white blood cell, RBC = red blood cell. **b,** JURKAT TALL cells were injected subcutaneously into immunocompromised mice. The following day, mice were given 10 mg/kg USP7i i.p. 3 times/week for 3 weeks. Shown is the mean tumor volume ± SD, measured using calipers (*n*=7 mice per treatment group, left panel). Weight of mice receiving JURKAT T-ALL cells via subcutaneous injection is also shown (*n*=7, right panel). **c,** Weight of mice receiving patient T-ALL cells upon treatment with USP7i (*n*=7 per treatment group). **d,** Mouse weight upon treatment with 10 mg/kg USP7i and 50 mg/kg GSKJ4 (vehicle, *n*=10; USP7i, *n*=6; USP7i+GSKJ4, *n*=10), following subcutaneous injection of JURKAT T-ALL cells.

**Supplementary Figure 13.**
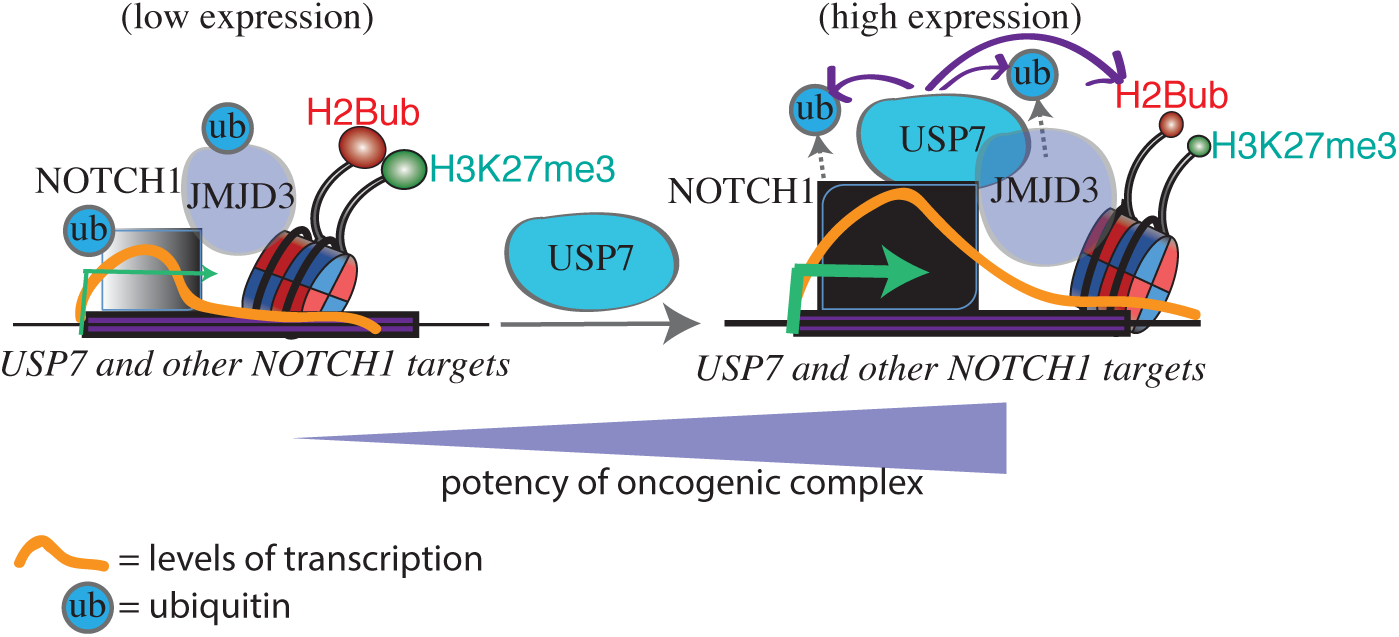
Model of the proposed action of USP7 in NOTCH1-driven T-ALL. The NOTCH1 complex positively controls expression of the *USP7* locus in T-ALL (left panel, *low expression).* Upon production, the USP7 protein is recruited by NOTCH1 to oncogenic targets, where it stabilizes NOTCH1/JMJD3 through deubiquitination (dotted arrows), leading to decreased H3K27me3 (through JMJD3 stabilization). USP7 also deubiquitinates H2B. Together, these events result is increased *(high)* expression of oncogenic targets. Curved arrows stemming from USP7 denote DUB activity on its substrates. Relative sizes of NOTCH1/JMJD3 and histone modifications represent relative amounts of proteins.

**Supplementary Table 1.** The JMJD3 interactome in T-ALL.

| Accession | Description | Avg Total Spectra |
| --- | --- | --- |
| O15054 | Lysine-specific demethylase 6B [KDM6B_HUMAN] | 184 |
| Q9UPU5 | Ubiquitin carboxyl-terminal hydrolase 24 [UBP24_HUMAN] | 43 |
| **B7Z815** | **Ubiquitin carboxyl-terminal hydrolase 7 [B7Z815_HUMAN]** | **33** |
| Q6KC79-2 | Isoform 2 of Nipped-B-like protein [NIPBL_HUMAN] | 22 |
| P31689 | DnaJ homolog subfamily A member 1 [DNJA1_HUMAN] | 20 |
| Q93008-1 | Isoform 2 of Probable ubiquitin carboxyl-terminal hydrolase FAF-X [USP9X_HUMAN] | 19 |
| Q6ZMT4 | Lysine-specific demethylase 7A [KDM7A_HUMAN] | 17 |
| Q96J02-2 | Isoform 2 of E3 ubiquitin-protein ligase Itchy homolog [ITCH_HUMAN] | 14 |
| Q9NNW5 | WD repeat-containing protein 6 [WDR6_HUMAN] | 13 |
| B3KTJ9 | cDNA FLJ38393 fis, clone FEBRA2007212 [B3KTJ9_HUMAN] | 13 |
| B4DNJ7 | Inosine-5'-monophosphate dehydrogenase [B4DNJ7_HUMAN] | 12 |
| P49915 | GMP synthase [glutamine-hydrolyzing] [GUAA_HUMAN] | 11 |
| Q8NFF5-3 | Isoform 3 of FAD synthase [FAD1_HUMAN] | 10 |
| Q92973-2 | Isoform 2 of Transportin-1 [TNPO1_HUMAN] | 10 |
| C7DJS2 | Glutathione S-transferase pi (Fragment) [C7DJS2_HUMAN] | 9 |
| Q5U5Z3 | Paraneoplastic antigen MA2 [Q5U5Z3_HUMAN] | 8 |
| O60884 | DnaJ homolog subfamily A member 2 [DNJA2_HUMAN] | 8 |
| O43823 | A-kinase anchor protein 8 [AKAP8_HUMAN] | 8 |
| P33176 | Kinesin-1 heavy chain [KINH_HUMAN] | 7 |
| E9PM69 | 26S protease regulatory subunit 6A [E9PM69_HUMAN] | 7 |
| B2RBA6 | cDNA, FLJ95407, highly similar to Homo sapiens MCM7 minichromosome maintenance deficient 7 (S. cerevisiae) (MCM7), mRNA [B2RBA6_HUMAN] | 7 |
| Q96K76-2 | Isoform 2 of Ubiquitin carboxyl-terminal hydrolase 47 [UBP47_HUMAN] | 7 |
| O43865 | Putative adenosylhomocysteinase 2 [SAHH2_HUMAN] | 7 |
| Q96D37 | Proto-oncogene vav [Q96D37_HUMAN] | 7 |
| Q9HBB9 | HC56 [Q9HBB9_HUMAN] | 6 |
| B7Z3G4 | cDNA FLJ60271, highly similar to SLIT-ROBO Rho GTPase-activating protein 2 [B7Z3G4_HUMAN] | 6 |
| B4DQ39 | cDNA FLJ55743, highly similar to DNA replication licensing factor MCM5 [B4DQ39_HUMAN] | 6 |
| P55345 | Protein arginine N-methyltransferase 2 [ANM2_HUMAN] | 6 |
| Q9BQA1 | Methylosome protein 50 [MEP50_HUMAN] | 6 |
| O75663 | TIP41-like protein [TIPRL_HUMAN] | 6 |
| Q9BTE3-2 | Isoform 2 of Mini-chromosome maintenance complex-binding protein [MCMBP_HUMAN] | 5 |
| B7Z254 | Protein disulfide-isomerase A6 [B7Z254_HUMAN] | 5 |
| B4DE69 | cDNA FLJ56053, highly similar to Serine/threonine-protein phosphatase 2A 65 kDa regulatory subunit A alpha isoform [B4DE69_HUMAN] | 5 |
| A8K5Y7 | cDNA FLJ78655, highly similar to Homo sapiens exportin 5 (XPO5), mRNA [A8K5Y7_HUMAN] | 5 |
| A8KA84 | cDNA FLJ78682, highly similar to Homo sapiens 2'-5'-oligoadenylate synthetase 3, 100kDa (OAS3), mRNA [A8KA84_HUMAN] | 5 |
| B7Z233 | Acetyl-CoA acetyltransferase, cytosolic [B7Z233_HUMAN] | 5 |
| P42677 | 40S ribosomal protein S27 [RS27_HUMAN] | 5 |
| Q9Y285 | Phenylalanine--tRNA ligase alpha subunit [SYFA_HUMAN] | 5 |
| F5GX11 | Proteasome subunit alpha type-1 [F5GX11_HUMAN] | 5 |
| P46109 | Crk-like protein [CRKL_HUMAN] | 5 |
| Q9Y266 | Nuclear migration protein nudC [NUDC_HUMAN] | 5 |
| Q7Z7L1 | Schlafen family member 11 [SLN11_HUMAN] | 5 |
| Q5HY54 | Filamin-A [Q5HY54_HUMAN] | 5 |
| B3KQ51 | cDNA FLJ32862 fis, clone TESTI2003598, highly similar to Serine/threonine-protein | 4 |
|  | phosphatase 2A catalytic subunit alpha isoform (EC 3.1.3.16) [B3KQ51_HUMAN] |  |
| G8JL95 | MAU2 chromatid cohesion factor homolog [G8JL95_HUMAN] | 4 |
| P61962 | DDB1- and CUL4-associated factor 7 [DCAF7_HUMAN] | 4 |
| P26640 | Valine--tRNA ligase [SYVC_HUMAN] | 4 |
| Q13185 | Chromobox protein homolog 3 [CBX3_HUMAN] | 4 |
| Q9NX07 | tRNA selenocysteine 1-associated protein 1 [TSAP1_HUMAN] | 4 |
| P06753-2 | Isoform 2 of Tropomyosin alpha-3 chain [TPM3_HUMAN] | 4 |
| B4DVR4 | cDNA FLJ60912, highly similar to Vinexin [B4DVR4_HUMAN] | 4 |
| Q9BSS9 | DNPEP protein (Fragment) [Q9BSS9_HUMAN] | 4 |
| Q14697 | Neutral alpha-glucosidase AB [GANAB_HUMAN] | 4 |
| Q9UJ70 | N-acetyl-D-glucosamine kinase [NAGK_HUMAN] | 3 |
| E9PPR1 | Elongation factor 1-delta (Fragment) [E9PPR1_HUMAN] | 3 |
| P62191 | 26S protease regulatory subunit 4 [PRS4_HUMAN] | 3 |
| P78406 | mRNA export factor [RAE1L_HUMAN] | 3 |
| B7Z7Z8 | Peptidyl-prolyl cis-trans isomerase FKBP5 [B7Z7Z8_HUMAN] | 3 |
| P23258 | Tubulin gamma-1 chain [TBG1_HUMAN] | 3 |
| P43686-2 | Isoform 2 of 26S protease regulatory subunit 6B [PRS6B_HUMAN] | 3 |
| Q8IYV2 | DEAD (Asp-Glu-Ala-Asp) box polypeptide 20 [Q8IYV2_HUMAN] | 3 |
| P62314 | Small nuclear ribonucleoprotein Sm D1 [SMD1_HUMAN] | 3 |
| P31146 | Coronin-1A [COR1A_HUMAN] | 3 |
| B4DV28 | cDNA FLJ54170, highly similar to Cytosolic nonspecific dipeptidase [B4DV28_HUMAN] | 3 |
| P49247 | Ribose-5-phosphate isomerase [RPIA_HUMAN] | 3 |
| Q15046 | Lysine--tRNA ligase [SYK_HUMAN] | 3 |
| C9JYI4 | Importin subunit alpha-5 (Fragment) [C9JYI4_HUMAN] | 3 |
| Q13085-3 | Isoform 3 of Acetyl-CoA carboxylase 1 [ACACA_HUMAN] | 3 |
| Q53H96 | Pyrroline-5-carboxylate reductase 3 [P5CR3_HUMAN] | 3 |
| B4E261 | cDNA FLJ55646, highly similar to Adapter-related protein complex 2 beta-1 subunit [B4E261_HUMAN] | 3 |
| B2R6S4 | cDNA, FLJ93089, highly similar to Homo sapiens NCK adaptor protein 1 (NCK1), mRNA [B2R6S4_HUMAN] | 3 |
| Q5JVF3-3 | Isoform 3 of PCI domain-containing protein 2 [PCID2_HUMAN] | 3 |
| Q9P2R3 | Ankyrin repeat and FYVE domain-containing protein 1 [ANFY1_HUMAN] | 3 |
| Q04695 | Keratin, type I cytoskeletal 17 [K1C17_HUMAN] | 3 |
| P46934-4 | Isoform 4 of E3 ubiquitin-protein ligase NEDD4 [NEDD4_HUMAN] | 3 |
| P22102-2 | Isoform Short of Trifunctional purine biosynthetic protein adenosine-3 [PUR2_HUMAN] | 3 |
| P62333 | 26S protease regulatory subunit 10B [PRS10_HUMAN] | 3 |
| B4DKL5 | cDNA FLJ51983, highly similar to Phosphoglycerate mutase 1 (EC 5.4.2.1) [B4DKL5_HUMAN] | 3 |
| B7ZVY5 | SLFN13 protein [B7ZVY5_HUMAN] | 3 |
| Q59FJ2 | Ubiquilin 1 isoform 1 variant (Fragment) [Q59FJ2_HUMAN] | 3 |
| B4E0B1 | cDNA FLJ52100 [B4E0B1_HUMAN] | 3 |
| B4DII5 | Importin subunit alpha [B4DII5_HUMAN] | 3 |
| Q71UM5 | 40S ribosomal protein S27-like [RS27L_HUMAN] | 3 |
| B7ZLC9 | GEMIN5 protein [B7ZLC9_HUMAN] | 3 |
| P51692 | Signal transducer and activator of transcription 5B [STA5B_HUMAN] | 3 |
| Q59GS3 | Proteasome 26S ATPase subunit 5 variant (Fragment) [Q59GS3_HUMAN] | 2 |
| Q08J23-2 | Isoform 2 of tRNA (cytosine(34)-C(5))-methyltransferase [NSUN2_HUMAN] | 2 |
| Q15181 | Inorganic pyrophosphatase [IPYR_HUMAN] | 2 |
| P36969-2 | Isoform Cytoplasmic of Phospholipid hydroperoxide glutathione peroxidase, mitochondrial [GPX4_HUMAN] | 2 |
| O14818-2 | Isoform 2 of Proteasome subunit alpha type-7 [PSA7_HUMAN] | 2 |
| Q86WI8 | Integrase interactor 1 protein isoform D (Fragment) [Q86WI8_HUMAN] | 2 |
| O60573 | Eukaryotic translation initiation factor 4E type 2 [IF4E2_HUMAN] | 2 |
| P61221 | ATP-binding cassette sub-family E member 1 [ABCE1_HUMAN] | 2 |
| O60547-2 | Isoform 2 of GDP-mannose 4,6 dehydratase [GMDS_HUMAN] | 2 |
| B7Z582 | Annexin [B7Z582_HUMAN] | 2 |
| H3BMM5 | Uncharacterized protein [H3BMM5_HUMAN] | 2 |
| P28066 | Proteasome subunit alpha type-5 [PSA5_HUMAN] | 2 |
| Q1RMY8 | GNB1 protein (Fragment) [Q1RMY8_HUMAN] | 2 |
| O00487 | 26S proteasome non-ATPase regulatory subunit 14 [PSDE_HUMAN] | 2 |
| P08579 | U2 small nuclear ribonucleoprotein B'' [RU2B_HUMAN] | 2 |
| Q8N395 | Putative uncharacterized protein DKFZp761N202 (Fragment) [Q8N395_HUMAN] | 2 |

## Methods

### Cell lines and primary cells

The human T-ALL cell lines CUTLL1 (gift from Iannis Aifantis, New York University), LOUCY (gift from Pieter Van Vlierberghe, Cancer Research Institute, Belgium), CCRF-CEM (American Type Culture Collection (ATCC), Manassas, VA, #CCL-119), JURKAT (ATCC) and KOPTK1 (Iannis Aifantis’ group) were cultured in RPMI 1640 medium supplemented with 10% heat-inactivated FBS (Sigma-Aldrich, St. Louis, MO), 2% penicillin/streptomycin (Gibco, Fisher Scientific, Hampton, NH), and 1% GlutaMAX (Gibco, Fisher Scientific). 293T cells (ATCC, #CRL-11268) were maintained in DMEM medium supplemented with 10% heat-inactivated FBS, 2% penicillin/streptomycin, and 1% GlutaMAX. Human pan T cells were purchased from AllCells.com (Alameda, CA). Primary human samples were collected by collaborating institutions with informed consent and analyzed under the supervision of the Institutional Review Board of Ghent University.

### Antibodies and reagents

The following antibodies were used: mouse anti-Actin (Millipore, Billerica, MA, clone C4), rabbit anti-JMJD3 (Cell Signaling Technology, Danvers, MA, #3457), rabbit anti-USP7 (Bethyl Laboratories, Montgomery, TX, #A300-033A-7), rabbit anti-Cleaved NOTCH1 (Val1744) (Cell Signaling Technology, #4147), rabbit anti-DNMT1 (Cell Signaling Technology, #5032), rabbit anti-H3K27Ac (Cell Signaling Technology, #8173S), rabbit anti-H3K27me3 (Cell Signaling Technology, #9733S), rabbit anti-H2BK120ub (Cell Signaling Technology, #5546S), rabbit anti-H2AK119ub (Cell Signaling Technology, #8240S), rabbit anti-H3K79me2 (gift from Ali Shilatifard, Northwestern University), rabbit anti-H3K4me3 (gift from Ali Shilatifard), and normal rabbit IgG (Millipore, #12-370). Secondary antibodies for Western blots were HRP-conjugated antirabbit and anti-mouse IgG (GE Healthcare, Chicago, IL).

Quick Start Bovine Gamma Globulin (BGG) Standard Set (protein standards) were purchased from Bio-Rad (Hercules, CA); benzonase, RNase A, dithiothreitol (DTT), EZview Red Anti-HA Affinity Gel (HA beads), and Influenza Hemagglutinin (HA) peptide were purchased from Sigma-Aldrich; NaV and NaF were purchased from New England BioLabs (Ipswich, MA); Protein G Dynabeads were purchased from Life Technologies (Carlsbad, CA); IgG-free BSA was purchased from Jackson ImmunoResearch Laboratories (West Grove, PA); phenol chloroform was purchased from ThermoScientific (Waltham, MA); and proteinase K, Tousimis formaldehyde, and Peptide International Z-Leu-Leu-Leu-H aldehyde (MG132) were purchased from Fisher Scientific. USP7i was a kind gift from Progenra (Malvern, PA). GSKJ4 was purchased from Cayman Chemical, Ann Arbor, MI, #12073)

### Immunoprecipitation (IP)

100 million T-ALL cells were collected and washed with chilled PBS. Cells were resuspended in 5 volumes of Buffer A (10 mM HEPES, 1.5 mM MgCl_2_, 10 mM KCl, 1:100 protease inhibitor (Sigma-Aldrich, P8340), 1 mM NaV, 1 mM NaF, and 0.5 mM DTT in H_2_O), incubated on ice for 10 min, and lysed using a Dounce homogenizer. Nuclear pellets were resuspended in 1.5 ml of TENT buffer (50 mM Tris pH 7.5, 5 mM EDTA, 150 mM NaCl, 0.05% v/v Tween 20, 1:100 protease inhibitor (Sigma-Aldrich, P8340), 1 mM NaV, 1 mM NaF, and 0.5 mM DTT in H_2_O) containing 5 mM MgCl_2_ and 100 units benzonase, and incubated at 4°C for 30 min, rotating. Lysates were passed through a 25^1/2^G needle/syringe 5 times, and spun down at 4°C, 2000 RPM, for 7 min to remove debris. Protein G magnetic beads were added to the lysates to decrease non-specific binding and incubated at 4°C for 30 min, rotating. Precleared lysates were then incubated with the appropriate antibody-conjugated beads (5 μg antibody per 100 million cells) at 4°C overnight, rotating. Beads were washed 4 times in TENT buffer at 4°C for 3 min, and protein complexes were eluted in 200 μl 0.1M glycine pH 2.5 for 10 min at 25°C, shaking. 20 μl of 1M Tris pH 8.0 was then added to the supernatants. For IP of HA-JMJD3, precleared lysates were incubated with HA-coated beads overnight, and protein complexes were eluted in 200 μl TENT buffer containing 400 μg/ml HA peptide (Sigma-Aldrich) at 4°C overnight, rotating.

### Mass spectrometry

Histone epiproteomics analysis was performed as previously described^1,2.^For the JMJD3 mass spectrometry the affinity-purified proteins were reduced, alkylated, and loaded onto an SDS-PAGE gel to remove any detergents and LCMS incompatible reagents. The gel plugs were excised, destained, and subjected to proteolytic digestion with trypsin. The resulting peptides were extracted and desalted as previously described^3^. An aliquot of the peptides was analyzed with LCMS coupled to a ThermoFisher Scientific Orbitrap QExactive Mass Spectrometer operated in data dependent mode. The data was searched against a UniProt human database, using Sequest within Proteome Discoverer. The results were filtered with a 1% FDR searched against a decoy database and for proteins with at least two unique peptides. Word clouds were generated using the statistical program R, and gene ontology was performed using the Gene Ontology Consortium (http://www.geneontology.org/).

### Western blot

To make total cell extracts, up to 10 million cells were collected and resuspended in 20 μl RIPA buffer (50 mM Tris HCl pH 8.0, 150 mM NaCl, 1% NP-40/IGEPAL, 0.5% sodium deoxycholate, 0.1% SDS, 1:100 protease inhibitor (Sigma-Aldrich, P8340), 1 mM NaV, and 1 mM NaF in H_2_O) per 1 million cells. Cells were lysed on ice for 20 min, and spun down at 4°C, max speed, for 10 min to remove debris.

Protein concentrations were determined via Bradford assay. Samples and buffer were diluted 1:10 in H_2_O. 2 μl of protein standards, H_2_O, or diluted sample were added to wells of a 96-well plate in duplicate. Then, 2 μl of diluted buffer and 100 μl Quick Start Bradford 1X Dye Reagent (Bio-Rad) were added to each well, and absorbance was measured at 600nm using the GloMax-Multi Detection System (Promega, Madison, WI).

Up to 50 μg sample was boiled in 1X SDS loading dye (Bio-Rad) at 95°C for 10 min prior to loading into 4-15% Tris-glycine polyacrylamide gels (Bio-Rad). 8 μl of PageRuler Plus Prestained Protein Ladder (10-250kD; Fisher Scientific) was also loaded. Gels were run at 100 V until samples reached the separating part of the gel, and then were run at 130V. Gels were transferred for 1.5h at 80V or overnight at 35-40V, and membranes were blocked in 5% milk in TBST (0.1% Tween 20 in 1X TBS) for 1h. Membranes were incubated at 4°C overnight with the appropriate antibody in TBST. Then, the membranes were washed 3 times for 10 min with TBST, incubated for 2h at 4°C with the appropriate secondary antibody, washed 3 times for 10 min with TBST, and developed using Clarity Western ECL Substrate (Bio-Rad), or SuperSignal West Femto Maximum Sensitivity Substrate (ThermoScientific) as needed, on a Bio-Rad ChemiDoc Touch Imaging System. Analysis was performed using Image Lab software (Bio-Rad).

### Chromatin immunoprécipitation (ChIP)

10 million T-ALL cells were cross-linked in 1 ml/million cells fixation buffer (1% formaldehyde, 1X PBS, and 1% FBS in H_2_O) for 10 min at 25°C. Then, 1:12.5 glycine [2.5M] was added for 5 min. Pelleted cells were then lysed according to the type of ChIP performed.

For histone ChIPs, cells were lysed in 375 μl of Nuclei Incubation Buffer (15 mM Tris pH 7.5, 60 mM KCl, 150 mM NaCl, 15 mM MgCl_2_, 1 mM CaCl_2_, 250 mM Sucrose, 0.3% NP-40, 1 mM NaV, 1 mM NaF, and 1 EDTA-free protease inhibitor tablet (Roche, Pleasanton, CA)/10 ml in H_2_O) for 10 min on ice. Nuclei were washed once with Digest Buffer (10 mM NaCl, 10 mM Tris pH 7.5, 3 mM MgCl_2_, 1 mM CaCl_2_, 1 mM NaV, 1 mM NaF, and 1 EDTA-free protease inhibitor tablet (Roche)/10 ml in H_2_O) and resuspended in 57 μl Digest Buffer containing 4.5 units MNase (USB, Cleveland, OH) for 1h at 37°C. MNase activity was quenched for 10 min on ice upon the addition of EDTA to a final concentration of 20 mM. Pelleted nuclei were lysed in 300 μl Nuclei Lysis Buffer (50 mM Tris-HCl pH 8.0, 10 mM EDTA pH 8.0, 1% SDS, 1 mM NaV, 1 mM NaF, and 1 EDTA-free protease inhibitor tablet (Roche)/10 ml in H_2_O) using a Bioruptor Pico (Diagenode, Denville, NJ) for 5 min (30 sec on, 30 sec off). Lysate was centrifuged at max speed for 5 min to remove debris, and 9 volumes of IP Dilution Buffer (0.01% SDS, 1.1% Triton X-100, 1.2 mM EDTA pH 8.0, 16.7 mM Tris-HCl pH 8.0, 167 mM NaCl, 1 mM NaV, 1 mM NaF, and 1 EDTA-free protease inhibitor tablet (Roche)/10 ml in H_2_O) were added to the supernatant. 50 μl protein G magnetic beads, blocked with IgG-free BSA, were added to the sample and incubated at 4°C for 30 min, rotating. 1% of the precleared sample was kept out as input, and the remaining sample was split into 3 tubes. 50 μl protein G magnetic beads conjugated to 15 μl of the appropriate antibody were added to each tube, and incubated at 4°C overnight, rotating. Bead-bound complexes were washed for 5 min each in 1 ml of Low Salt Buffer (20 mM Tris-HCl pH 8.0, 150 mM NaCl, 2 mM EDTA, 1% w/v Triton X-100, and 0.1% w/v SDS in H_2_O), High Salt Buffer (20 mM Tris-HCl pH 8.0, 500 mM NaCl, 2 mM EDTA, 1% w/v Triton X-100, and 0.1% w/v SDS in H_2_O), LiCl Buffer (10 mM Tris-HCl pH8.0, 250 mM LiCl, 1 mM EDTA, 1% w/v NP-40, and 1% w/v deoxycholic acid in H_2_O), and TE.

For epigenetic regulator and transcription factor ChIPs, cells were lysed in 1 ml LB1 Buffer (50 mM HEPES-KOH pH 7.5, 140 mM NaCl, 1 mM EDTA, 10% Glycerol, 0.5% NP-40, 0.25% Triton X-100, 1:100 protease inhibitor (Sigma-Aldrich, P8340), 1 mM NaV, and 1 mM NaF in H_2_O) for 10 min at 4°C, rotating. Nuclei pellets were resuspended in LB2 Buffer (10 mM Tris-HCl pH 8.0, 200 mM NaCl, 1 mM EDTA, 0.5 mM EGTA, 1:100 protease inhibitor (Sigma-Aldrich, P8340), 1 mM NaV, and 1 mM NaF in H_2_O) and incubated for

10 min at 25°C, rotating. Finally, nuclei pellets were lysed in 300 μl LB3 Buffer (10 mM Tris-HCl pH 8.0, 100 mM NaCl, 1 mM EDTA pH 8.0, 0.5 mM EGTA pH 8.0, 0.1% sodium deoxycholate, 0.5% N-lauroylsarcosine, 1:100 protease inhibitor (Sigma-Aldrich, P8340), 1 mM NaV, and 1 mM NaF in H_2_O) using a Bioruptor Pico (Diagenode) for 5 min (30 sec on, 30 sec off). 1% Triton X-100 was added, and samples were centrifuged at max speed to remove debris. 50 μl protein G magnetic beads, blocked with IgG-free BSA, were added to the sample and incubated at 4°C for 30 min, rotating. 1% of the precleared sample was kept out as input, and the remaining sample was used for IP. 50 μl protein G magnetic beads conjugated to 5-10 μg of the appropriate antibody were added to each tube, and incubated at 4°C overnight, rotating. Bead-bound complexes were washed for 5 min each in 1 ml of RIPA Wash Buffer (50 mM HEPES-KOH pH 7.6, 300 mM LiCl, 1 mM EDTA, 1% NP-40, and 0.7% sodium deoxycholate in H_2_O), 5 times, and 1 time in 1 ml TE Wash Buffer (10 mM Tris-HCl pH 8.0, 1 mM EDTA, and 50 mM NaCl in H_2_O).

To elute bead-bound complexes, 50 μl of Elution Buffer (for 10 ml total volume, 7.5 ml H_2_O, 0.5 ml 20% SDS, and 2 ml 0.5M sodium bicarbonate) was added to each sample, and samples were incubated at 65°C for 15 min, shaking at 1000 RPM on a thermomixer (ThermoScientific). Elution was repeated a second time, and then 100 μl RNase Buffer (12 μl of 5M NaCl, 0.2 μl 30 mg/ml RNase, and 88 μl TE) was added to each ChIP and input sample. Samples were incubated at 37°C for 20 min, followed by the addition of PK Buffer (2.5 μl 20 mg/ml proteinase K, 5 μl 20% SDS, and 92.5 μl TE) overnight at 65°C. An equal volume of phenol chloroform solution was added to the samples, which were vortexed for 1 min and transferred to MaXtract High Density tubes (Qiagen). Samples were centrifuged for 8 min at max speed, and the upper phase was transferred to new tubes containing 1.5 μl 20 mg/ml glycogen. Then, 30 μl sodium acetate and 800 μl 100% ethanol were added, and tubes were incubated on dry ice to 30-60 min. DNA pellets were washed in 70% ethanol, air-dried, and resuspended in 30 μl H_2_O.

### ChIP-Seq

Libraries were prepared using Agencourt AMPure XP beads (Beckman Coulter, Brea, CA) and the KAPA HTP Library Preparation Kit (KAPA Biosystems, Wilmington, MA), according to the manufacturer’s protocol (v4.15). DNA fragment size was determined using High Sensitivity DNA Chips read on a 2100 Bioanalyzer (Agilent Technologies, Santa Clara, CA). Libraries were sequenced on the Illumina NextSeq 500 (San Diego, CA; 50bp single reads). FASTQ reads were first trimmed from the 3’ end using Trimmomatic^4^ version 0.33, such that the Phred33 score of all the nucleotides was above 30 and all reads shorter than 20bp were discarded. The resulting reads were then aligned to the hg19 genome using bowtie version 1.1.2 with the following parameters: bowtie ‐p 5 ‐m 1 ‐v 2 ‐S^5^. Next, the bam file was sorted and bigwig tracks were created by extending each read by 150 bp and using the GenomicAlignments^6^ package in R in order to calculate the coverage of reads in counts per million (CPM) normalized to the total number of reads for each sample in the library.

### Peak calling

Peaks were called using MACS^7^ version 1.4.2 with the default parameters, using the specific aligned ChIP-Seq data for a particular antibody as the treatment sample and an aligned bam file of the unprecipitated input as a control sample. Only the top 10,000 peaks ordered by peak score (which is proportional to the FDR of the peak) were chosen for further analysis.

### Calculation of overlaps and statistical significance

Overlap between two ChIP-Seq peak datasets were calculated using the mergePeaks tool in the HOMER^8^ tools. In brief, the intersection between two ChIP-Seq peak datasets was determined by calculating the number of peaks in one set that overlap with the peaks in the second set. Statistical significance of the overlaps between ChIP-Seq datasets was done by using reservoir sampling^9^ with a set of 161 transcription factor peak binding sites downloaded from http://hgdownload.cse.ucsc.edu/goldenpath/hg19/encodeDCC/wgEncodeRegTfbsClustered/ (wgEncodeRegTfbsClusteredV3.bed.gz, ENCODE data). The tool for implementing the sampling schema is called poverlaps and can be downloaded from github at https://github.com/brentp/poverlap. A total of 100 permutations per pair-wise combination of transcription factor overlaps was used. Venn diagrams of overlaps were generated using an online Venn diagram generator (http://jura.wi.mit.edu/bioc/tools/venn.php).

### Motif analysis and pathway enrichment analysis

HOMER, with the standard default parameters, was used to determine the enrichment for known and unknown motifs in the USP7 ChIP-Seq peaks. Pathway enrichment analysis of USP7-bound genes were determined using GREAT (Genomic Regions Enrichment of Annotations Tool)^10^.

### Generation of heatmaps

Heatmap representation of log2 fold-changes in epigenetic marks between DMSO and USP7 inhibition were visualized using ngs.plot^11^. In brief, log2 fold-changes of H3K27me3, H2Aub, H2Bub, and H3K79me2 upon USP7 inhibition versus DMSO treatment at the top 10,000 USP7 peaks were visualized. A k-means clustering algorithm option was used with the number of clusters set to 5. Correlated gene expression changes were calculated by determining the log2 fold-change in gene expression upon USP7 or JMJD3 inhibition of the nearest protein-coding gene to the USP7 peak. Gene expression changes were visualized using Java TreeView^12^.

### Quantitative polymerase chain reaction (qPCR)

RNA was isolated from T-ALL cells using the Aurum Total RNA Mini Kit (Bio-Rad), and was quantified on a Qubit 3.0 Fluorometer (Life Technologies) using the Qubit RNA HS (High Sensitivity) Assay Kit (Life Technotogies) according to the manufacturers’ instructions. cDNA was made using the High Capacity cDNA RT Kit (Applied Biosystems, Foster City, CA), according to the manufacturer’s protocol. qPCR reactions were carried out using iTaq Universal SYBR Green Supermix (Bio-Rad) and the following primers: USP7 – 5’-CGGTGTTGTGTCCATCACTC-3’ (forward) and 5’ ‐AGTTGAGCGAGCCCGAG-3’ (reverse), NOTCH3 – 5’ ‐GTAGAGGGCATGGTGGAAGA-3’ (forward) and 5’ ‐AAGTGGTCCAACAGCAGCTT-3’ (reverse), DTX1 – 5’ ‐CTCGCCACTGCTATCT ACCC-3’ (forward) and 5’ ‐CGTGCCGATAGTGAAGATGA-3’ (reverse), and HES1 – 5’ ‐GCAGATGACGGCTGCGCTGA-3’ (forward) and 5’-AAGCGGGTCACCTCGTTCATGC-3’ (reverse). Samples were run on a CFX Connect Reak-Time System (Bio-Rad) under the following conditions: 95°C for 10 sec (denaturation), 60°C for 30 sec (annealing), and 72°C for 30 sec (elongation), for 40 cycles. Analyses were performed using GraphPad Prism software (GraphPad Software, La Jolla, CA). Statistical comparisons were made using the Student’s unpaired, two-sided t-test. p values <0.05 were considered statistically significant. Relative mRNA abundance refers to the levels of the gene of interest normalized to housekeeping genes GAPDH or G6PD. All values were then normalized to the control sample.

### Analysis of data from publically available databases

Analysis of microarray data from GEO was done using the NCBI GEO2R online tool for microarray analysis. Adjusted p-value calculations were done using Benjamini & Hochberg (False discovery rate or FDR) option. A FDR of <0.05 was considered to be statistically significant.

### Generation of CRISPR cell lines

Single-guide RNAs (sgRNAs) targeting USP7 (sgRNA 1, 5’ ‐GAGTGATGGACACAACACCGCGG-3’; sgRNA 2, 5’-AGACACCAGTTGGCGCTCCGAGG-3’), designed using CHOPCHOP (http://chopchop.cbu.uib.no), were cloned into pLX-sgRNA (Addgene, Cambridge, MA, #50662). pLX-sgRNA was amplified using F1 (5’-AAACTCGAGTGTACAAAAAAGCAGGCTTTAAAG-3’) and R1 (sgRNA 1, 5’-CGGTGTTGTGTCCATCACTCGGTGTTTCGTCCTTTCC-3’; sgRNA 2, 5’-GGAGCGCCAACTGGTGTCTCGGTGTTTCGTCCTTTCC-3’), and F2 (sgRNA 1, 5’-GAGTGATGGACACAACACCGGTTTTAGAGCTAGAAATAGCAA-3’; sgRNA 2, 5’-GAGACACCAGTTGGCGCTCCGTTTTAGAGCTAGAAATAGCAA-3’) and R2 (5’-AAAGCTAGCTAATGCCAACTTTGTACAAGAAAGCTG-3’). These products were gel-purified using the QIAquick Gel Extraction Kit (Qiagen, Germantown, MD), and amplified using F1 and R2. PCR products were digested with NheI and XhoI, ligated, and used for STBL3 (ThermoScientific) bacterial transformation, according to the manufacturer’s protocol.

To make virus, 293T cells were seeded in a 6-well plate. The next day, 0.25 M CaCl_2_, 1X HBS pH 7.05, 2 μg VSVG envelope plasmid, and 3 μg pCMV-dR8.91 (Delta 8.9) lentiviral packaging plasmid were added to 10 μg DNA (sgRNA or empty vector) and added dropwise to the cells. Medium was changed the next day, and after 24h, virus was collected 3 times (6-8h between collections). Pooled viral supernatant (3 ml) was filtered using a 0.45 μM syringe filter, and 1 ml viral supernatant was added to a 6-well plate containing 0.5 million JURKAT T-ALL cells stably expressing pCW-Cas9 (Addgene, #83481), a gift from Navdeep Chandel (Northwestern University). Infection was repeated the next day, and 1 week later, cells were treated for 3 weeks with 5 μg/ml blasticidin to select for cells expressing the sgRNA or empty vector. 1 μg/ml doxycycline was then added to cells to induce the disruption of USP7.

### Ubiquitin competition assay

This assay was performed as previously described, using recombinant enzymes and a panel of USP7 inhibitors, including P217564^13^.

### Knockdown of USP7 by shRNA

The following USP7-specific shRNA (Sigma-Aldrich) was used: 5'-CCGGCCTGGATTTGTGGTTACGTTACTCGAGTAACGTAACCACAAATCCAGGTTTTT-3'. The shRNA was cloned into puro-IRES-GFP, and retrovirus was used to infect JURKAT T-ALL cells as previously described^14^.

### Viability assays, apoptosis, and cell cycle analysis

500,000 cells were plated in wells of a 24-well plate, and treated with DMSO or USP7i as indicated in the Figure Legends. Cells were counted each day via trypan blue, and inhibitor/medium was changed. When cells reached a confluency of >1 million cells/ml, cells were diluted 1:2, and this dilution was factored into the cell numbers for viability assays. For apoptosis analysis, cells were stained with LIVE/DEAD Fixable Near-IR Dead CeU Stain (Life Technotogies) according to the manufacturer’s protocol except that cells were stained for 20 min at 4°C, prior to staining with PE-conjugated Annexin V (Life Technologies) in Annexin V Binding Buffer (BD Biosciences, San Jose, CA), according to the manufacturers’ instructions. To measure cell cyde, cells were fixed in 100 μl Fix and Perm Medium A (Life Technologies) for 15 min, washed with PBS, and incubated with 100 μl Fix and Perm Medium B (Life Technologies) + 1 μg/ml DAPI (Invitrogen, Carlsbad, CA) for 1h at 4°C. Flow cytometry was performed on an LSR II (BD, Franklin Lakes, NJ) and analyses were performed using FlowJo software (Tree Star, Ashland, OR). Statistical analyses were performed using GraphPad Prism software (GraphPad Software). Comparisons were made using the Student’s unpaired, two-sided t-test. p values <0.05 were considered statistically significant.

### TUBE ubiquitin assay

200 million JURKAT T-ALL cells were treated in duplicate with either DMSO or 5 μM USP7i for 8h. Cell pellets were collected, washed with PBS, and frozen. Ubiquitinated substrates were pulled down using agarose-TUBE beads, eluted, and treated with the deubiquitinase USP2 to remove ubiquitin prior to Western blotting as previously described^13^.

### RNA-Seq

RNA was extracted from up to 10 million T-ALL cells using the Aurum Total RNA Mini Kit (Bio-Rad) according to the manufacturer’s instructions. RNA was quantified on a Qubit 3.0 Fluorometer (Life Technologies) using the Qubit RNA HS (High Sensitivity) Assay Kit (Life Technologies) according to the manufacturer’s instructions. RNA integrity and DNA fragment size were determined using RNA Nano Chips and High Sensitivity DNA Chips read on a 2100 Bioanalyzer (Agilent Technologies). Libraries were prepared using Agencourt AMPure XP beads (Beckman Coulter) and the TruSeq RNA Library Prep Kit v2 (Illumina), according to the manufacturer’s Low Sample (LS) protocol and sequenced on the Illumina NextSeq 500 (50bp single reads). FASTQ reads were aligned to the hg19 genome using tophat version 2.1.0^15^ using the following options ‐‐no-novel-juncs ‐‐read-mismatches 2 ‐‐read-edit-dist 2 ‐‐max-multihits 5. The generated bam files were then used to count the reads only at the exons of genes using htseq-count^16^ with the following parameters ‐q ‐m intersection-nonempty ‐s reverse ‐t exon. Differential expression analysis was done using the R package edgeR^17^. Bigwig tracks of RNA-Seq expression were generated by using the GenomicAlignments package in R in order to calculate the coverage of reads in counts per million (CPM) normalized to the total number of uniquely mapped reads for each sample in the library.

### Superenhancer analysis

Superenhancers were determined using a H3K27Ac ChIP-Seq dataset in JURKAT cells treated with DMSO or USP7 inhibitor. Aligned bam files of H3K27Ac ChIP-Seq signal and input control were analyzed using the Rank Ordering of Super-Enhancers algorithm (ROSE)^18,19^. A stitching distance of 12.5 kbp were used to stich together enhancer regions, and regions within 2.5 kbp of a transcriptional start site were ignored in order to prevent promoter bias. The initial set of enhancer regions used in the analysis was determined by merging the peaks of H3K4me1 and H3K27Ac in CUTLL1 cells (ChIP-Seq dataset for H3K4me1 signal and H3K27Ac was downloaded from GEO with the accession numbers GSM732910 and GSM1252938 for H3K4me1 and H3K27Ac, respectively).

### Gene set enrichment analysis

Gene set enrichment analysis (GSEA) was done using the Broad Institute GSEA software (http://software.broadinstitute.org/gsea/index.jsp)^20,21^. RNA-Seq gene expression data were used to create a preranked order of genes, where genes were sorted using the p-value of the differential expression between USP7 inhibition and DMSO treatment, and gene up or downregulation upon USP7 inhibition was determined. In brief, the genes were ranked by calculating ‐log(p-value)*sign(logFC), and a PreRanked GSEA analysis was done using the Hallmark database and the standard weighted enrichment statistic with 5,000 permutations. Enriched datasets with a FDR < 0.05 were considered statistically significant.

### Inhibitor wash-off experiment

45 million JURKAT T-ALL cells were treated with DMSO, 1 μM USP7i, or 1 μM USP7i + 2 μM GSKJ4 for 24h, with medium and inhibitors replaced at 12h. 5 million cells from each flask were collected for RNA extraction, and the remaining 40 million cells in each flask were divided equally and treated with either DMSO or GSKJ4 for 2h, 4h, 6h, or overnight, and cells were collected at each time point for RNA extraction.

### Intravenous and subcutaneous xenograft studies

All mice were housed in a barrier facility, and procedures were performed as approved by the Northwestern University Institutional Animal Care and Use Committee (protocol Ntziachristos #IS00002058 and Mazar #IS00000556) and the Ghent University Animal Ethics Committee (protocol #ECD16/56).

For JURKAT T-ALL subcutaneous studies, 0.7 million cells were injected subcutaneously into the right flank of 8-week-old NOD.Cg-Prkdcscid male mice (#005557, Jackson Laboratories, Portage, MI) with an equal volume of BD Matrigel, at 50 μl of cells to 50 μl Matrigel. The following day, mice were randomly assigned to treatment groups (7 mice/group), and were treated with either DMSO or 10 mg/kg USP7i 3 times per week i.p. for 8 doses. One week later, mice were treated 5 times per week i.p. for 17 days. Body weight and tumor size (via calipers) were measured 3 times per week.

For JURKAT T-ALL transplant studies, 1 million cells were injected via tail vein into 8-week-old NOD.Cg-Prkdcscid male mice (#005557, Jackson Laboratories) in 100 μl PBS. Animals were monitored by IVIS every 3 days until luciferase signal was detected, and then animals were randomly assigned to treatment groups (9 mice/group). Mice were treated 3 times per week i.p. with DMSO or 10 mg/kg USP7i (in 10% DMSO + 2% Tween 80) for up to 10 doses. IVIS images were taken twice per week (Perkin Elmer) and animal weight was measured 3 times per week to ensure accurate dosing.

For primagraft studies, 1.2 million primary human T-ALL cells (from the spleen of leukemic primary xenograft engrafted with cells of T-ALL patient FV, N16/0267) in 150 μl PBS were injected via tail vein into 1.5-month-old female NGS mice (Jackson Laboratories). Two weeks post-transplant, blood was collected via tail nick for analysis of hCD45 expression (engraftment) using PE-conjugated mouse anti-human CD45 antibody (Miltenyi Biotec, Auburn, CA; clone 5B1). Flow cytometry was performed on and analyzed using software FlowJo. One week later, mice were randomly divided into two groups (7 mice/group) and treated with either DMSO or 10 mg/kg USP7i in 2% Tween 80 + 10% DMSO in PBS. Treatment was administered on days 1-5 and 8-11 i.v., and on days 12 and 15-22 i.p. Blood was collected via tail nick on days 17 and 22 to measure tumor burden by staining for hCD45.

For toxicity studies, 8-week-old NOD.Cg-Prkdcscid male mice (#005557, Jackson Laboratories) were randomly assigned to treatment groups (3 mice/group). Mice were treated with either vehicle (10% DMSO, 2% Tween-80, and 10% captisol in PBS), 10 mg/kg USP7i, 10 mg/kg USP7i + 10 mg/kg GSKJ4, 10 mg/kg USP7i + 20 mg/kg GSKJ4, or 10 mg/kg USP7i + 50 mg/kg GSKJ4 i.v. daily for 5 days to evaluate treatment toxicity.

For combination treatment studies, 1 million JURKAT T-ALL cells were injected subcutaneously into the right flank of 8-week-old NOD.Cg-Prkdcscid male mice (#005557, Jackson Laboratories) with an equal volume of BD Matrigel, at 50 μl of cells to 50 μl Matrigel. After tumors reached 150-200 mm^3^, mice were randomly assigned to treatment groups and treated i.v. with vehicle (10% DMSO, 2% Tween 80, and 10% captisol in PBS; *n*=10), 10 mg/kg USP7i (*n*=6 per inhibitor), or 10 mg/kg USP7i + 50 mg/kg GSKJ4 (*n*=10) 5 times the first week, and then 3 times per week until tumors reached 1.5 cm^3^.

Analyses were performed using GraphPad Prism software (GraphPad Software). Statistical comparisons were made using the Student’s unpaired, two-sided t-test. p values <0.05 were considered statistically significant.

### Complete blood cell count analysis

Mouse blood samples were collected via cardiac puncture and run on a Hemavet 950 FS (Drew Scientific, Inc., Miami Lakes, FL) to obtain blood cell counts.

## References

1 Hunger, S. P. & Mullighan, C. G. Acute Lymphoblastic Leukemia in Children. The New England journal of medicine 373, 1541–1552, doi:10.1056/NEJMra1400972 (2015).

2 Pui, C. H., Relling, M. V. & Downing, J. R. Acute lymphoblastic leukemia. The New England journal of medicine 350, 1535–1548, doi:10.1056/NEJMra023001 (2004).

3 Pui, C. H., Mullighan, C. G., Evans, W. E. & Relling, M. V. Pediatric acute lymphoblastic leukemia: where are we going and how do we get there? Blood 120, 1165–1174, doi:10.1182/blood-2012-05-378943 (2012).

4 Pui, C. H. & Evans, W. E. Treatment of acute lymphoblastic leukemia. The New England journal of medicine 354, 166–178, doi:10.1056/NEJMra052603 (2006).

5 Aifantis, I., Raetz, E. & Buonamici, S. Molecular pathogenesis of T-cell leukaemia and lymphoma. Nat Rev Immunol 8, 380–390, doi:10.1038/nri2304 (2008).

6 Pui, C. H., Robison, L. L. & Look, A. T. Acute lymphoblastic leukaemia. Lancet 371, 1030–1043, doi:10.1016/S0140-6736(08)60457-2 (2008).

7 Haydu, J. E. & Ferrando, A. A. Early T-cell precursor acute lymphoblastic leukaemia. Curr Opin Hematol 20, 369–373, doi:10.1097/MOH.0b013e3283623c61 (2013).

8 Coustan-Smith, E. et al. Early T-cell precursor leukaemia: a subtype of very high-risk acute lymphoblastic leukaemia. Lancet Oncol 10, 147–156, doi:10.1016/S1470-2045(08)70314-0 (2009).

9 Weng, A. P. et al. Activating mutations of NOTCH1 in human T cell acute lymphoblastic leukemia. Science 306, 269–271, doi:10.1126/science.1102160 (2004).

10 Real, P. J. et al. Gamma-secretase inhibitors reverse glucocorticoid resistance in T cell acute lymphoblastic leukemia. Nature medicine 15, 50–58, doi:10.1038/nm.1900 (2009).

11 Tosello, V. et al. WT1 mutations in T-ALL. Blood 114, 1038–1045, doi:10.1182/blood-2008-12-192039 (2009).

12 De Keersmaecker, K. & Ferrando, A. A. TLX1-induced T-cell acute lymphoblastic leukemia. Clin Cancer Res 17, 6381–6386, doi:10.1158/1078-0432.CCR-10-3037 (2011).

13 Zenatti, P. P. et al. Oncogenic IL7R gain-of-function mutations in childhood T-cell acute lymphoblastic leukemia. Nature genetics 43, 932–939, doi:10.1038/ng.924 (2011).

14 Ntziachristos, P. et al. Genetic inactivation of the polycomb repressive complex 2 in T cell acute lymphoblastic leukemia. Nature medicine 18, 298–301, doi:10.1038/nm.2651 (2012).

15 Ntziachristos, P. et al. Contrasting roles of histone 3 lysine 27 demethylases in acute lymphoblastic leukaemia. Nature, doi:10.1038/nature13605 (2014).

16 Trimarchi, T. et al. Genome-wide mapping and characterization of Notch-regulated long noncoding RNAs in acute leukemia. Cell 158, 593–606, doi:10.1016/j.cell.2014.05.049 (2014).

17 Buonamici, S. et al. CCR7 signalling as an essential regulator of CNS infiltration in T-cell leukaemia. Nature 459, 1000–1004, doi:10.1038/nature08020 (2009).

18 Mullighan, C. G. & Downing, J. R. Global genomic characterization of acute lymphoblastic leukemia. Seminars in hematology 46, 3–15, doi:10.1053/j.seminhematol.2008.09.005 (2009).

19 Zhang, J. et al. The genetic basis of early T-cell precursor acute lymphoblastic leukaemia. Nature 481, 157–163, doi:10.1038/nature 10725 (2012).

20 Roberts, K. G. & Mullighan, C. G. Genomics in acute lymphoblastic leukaemia: insights and treatment implications. Nat Rev Clin Oncol 12, 344–357, doi:10.1038/nrclinonc.2015.38 (2015).

21 Lecona, E., Narendra, V. & Reinberg, D. USP7 cooperates with SCML2 to regulate the activity of PRC1. Molecular and cellular biology 35, 1157–1168, doi:10.1128/MCB.01197-14 (2015).

22 Zhang, Z. M. et al. An Allosteric Interaction Links USP7 to Deubiquitination and Chromatin Targeting of UHRF1. Cell reports 12, 1400–1406, doi:10.1016/j.celrep.2015.07.046 (2015).

23 Song, M. S. et al. The deubiquitinylation and localization of PTEN are regulated by a HAUSP-PML network. Nature 455, 813–817, doi:10.1038/nature07290 (2008).

24 Li, M., Brooks, C. L., Kon, N. & Gu, W. A dynamic role of HAUSP in the p53-Mdm2 pathway. Mol Cell 13, 879–886 (2004).

25 Li, M. et al. Deubiquitination of p53 by HAUSP is an important pathway for p53 stabilization. Nature 416, 648–653, doi:10.1038/nature737 (2002).

26 Kon, N. et al. Inactivation of HAUSP in vivo modulates p53 function. Oncogene 29, 1270–1279, doi:10.1038/onc.2009.427 (2010).

27 van der Horst, A. et al. FOXO4 transcriptional activity is regulated by monoubiquitination and USP7/HAUSP. Nat Cell Biol 8, 1064–1073, doi:10.1038/ncb1469 (2006).

28 Du, Z. et al. DNMT1 stability is regulated by proteins coordinating deubiquitination and acetylation-driven ubiquitination. Sci Signal 3, ra80, doi:10.1126/scisignal.2001462 (2010).

29 Espinosa, J. M. Histone H2B ubiquitination: the cancer connection. Genes & development 22, 2743–2749, doi:10.1101/gad.1732108 (2008).

30 Chen, S. T. et al. The Deubiquitinating Enzyme USP7 Regulates Androgen Receptor Activity by Modulating Its Binding to Chromatin. J Biol Chem 290, 21713–21723, doi:10.1074/jbc.M114.628255 (2015).

31 Sowa, M. E., Bennett, E. J., Gygi, S. P. & Harper, J. W. Defining the human deubiquitinating enzyme interaction landscape. Cell 138, 389–403, doi:10.1016/j.cell.2009.04.042 (2009).

32 Liefke, R., Karwacki-Neisius, V. & Shi, Y. EPOP Interacts with Elongin BC and USP7 to Modulate the Chromatin Landscape. Mol Cell 64, 659–672, doi:10.1016/j.molcel.2016.10.019 (2016).

33 Chauhan, D. et al. A small molecule inhibitor of ubiquitin-specific protease-7 induces apoptosis in multiple myeloma cells and overcomes bortezomib resistance. Cancer Cell 22, 345–358, doi:10.1016/j.ccr.2012.08.007 (2012).

34 Agger, K. et al. UTX and JMJD3 are histone H3K27 demethylases involved in HOX gene regulation and development. Nature 449, 731–734, doi:10.1038/nature06145 (2007).

35 Agger, K. et al. The H3K27me3 demethylase JMJD3 contributes to the activation of the INK4A-ARF locus in response to oncogene‐ and stress-induced senescence. Genes & development 23, 1171–1176, doi:10.1101/gad.510809 (2009).

36 De Santa, F. et al. The histone H3 lysine-27 demethylase Jmjd3 links inflammation to inhibition of polycomb-mediated gene silencing. Cell 130, 1083–1094, doi:10.1016/j.cell.2007.08.019 (2007).

37 De Santa, F. et al. Jmjd3 contributes to the control of gene expression in LPS-activated macrophages. The EMBO journal 28, 3341–3352, doi:10.1038/emboj.2009.271 (2009).

38 Thompson, B. J. et al. The SCFFBW7 ubiquitin ligase complex as a tumor suppressor in T cell leukemia. J Exp Med 204, 1825–1835, doi:10.1084/jem.20070872 (2007).

39 Thompson, B. J. et al. Control of hematopoietic stem cell quiescence by the E3 ubiquitin ligase Fbw7. J Exp Med 205, 1395–1408, doi:10.1084/jem.20080277 (2008).

40 Skaar, J. R., Pagan, J. K. & Pagano, M. SCF ubiquitin ligase-targeted therapies. Nat Rev Drug Discov 13, 889–903, doi:10.1038/nrd4432 (2014).

41 Nijman, S. M. et al. A genomic and functional inventory of deubiquitinating enzymes. Cell 123, 773–786, doi:10.1016/j.cell.2005.11.007 (2005).

42 Komander, D., Clague, M. J. & Urbe, S. Breaking the chains: structure and function of the deubiquitinases. Nat Rev Mol Cell Biol 10, 550–563, doi:10.1038/nrm2731 (2009).

43 Yatim, A. et al. NOTCH1 nuclear interactome reveals key regulators of its transcriptional activity and oncogenic function. Mol Cell 48, 445–458, doi:10.1016/j.molcel.2012.08.022 (2012).

44 Fan, Y. H. et al. USP7 inhibitor P22077 inhibits neuroblastoma growth via inducing p53-mediated apoptosis. Cell Death Dis 4, e867, doi:10.1038/cddis.2013.400 (2013).

45 Wang, L. et al. Ubiquitin-specific Protease-7 Inhibition Impairs Tip60-dependent Foxp3+ T-regulatory Cell Function and Promotes Antitumor Immunity. EBioMedicine 13, 99–112, doi:10.1016/j.ebiom.2016.10.018 (2016).

46 Hjerpe, R. et al. Efficient protection and isolation of ubiquitylated proteins using tandem ubiquitin-binding entities. EMBO Rep 10, 1250–1258, doi:10.1038/embor.2009.192 (2009).

47 Herranz, D. et al. A NOTCH1-driven MYC enhancer promotes T cell development, transformation and acute lymphoblastic leukemia. Nature medicine, doi:10.1038/nm.3665 (2014).

48 Wang, H. et al. NOTCH1-RBPJ complexes drive target gene expression through dynamic interactions with superenhancers. Proceedings of the National Academy of Sciences of the United States of America 111, 705–710, doi:10.1073/pnas.1315023111 (2014).

49 Hnisz, D. et al. Super-enhancers in the control of cell identity and disease. Cell 155, 934–947, doi:10.1016/j.cell.2013.09.053 (2013).

50 Whyte, W. A. et al. Master transcription factors and mediator establish super-enhancers at key cell identity genes. Cell 153, 307–319, doi:10.1016/j.cell.2013.03.035 (2013).

51 Filippakopoulos, P. et al. Selective inhibition of BET bromodomains. Nature 468, 1067–1073, doi:10.1038/nature09504 (2010).

52 Delmore, J. E. et al. BET bromodomain inhibition as a therapeutic strategy to target c-Myc. Cell 146, 904–917, doi:10.1016/j.cell.2011.08.017 (2011).

53 van der Knaap, J. A. et al. GMP synthetase stimulates histone H2B deubiquitylation by the epigenetic silencer USP7. Mol Cell 17, 695–707, doi:10.1016/j.molcel.2005.02.013 (2005).

54 van der Knaap, J. A., Kozhevnikova, E., Langenberg, K., Moshkin, Y. M. & Verrijzer, C. P. Biosynthetic enzyme GMP synthetase cooperates with ubiquitin-specific protease 7 in transcriptional regulation of ecdysteroid target genes. Molecular and cellular biology 30, 736–744, doi:10.1128/MCB.01121-09 (2010).

55 Strikoudis, A. et al. Regulation of transcriptional elongation in pluripotency and cell differentiation by the PHD-finger protein Phf5a. Nat Cell Biol 18, 1127–1138, doi:10.1038/ncb3424 (2016).

56 Strikoudis, A., Lazaris, C., Ntziachristos, P., Tsirigos, A. & Aifantis, I. Opposing functions of H2BK120 ubiquitylation and H3K79 methylation in the regulation of pluripotency by the Paf1 complex. Cell cycle, 0, doi:10.1080/15384101.2017.1295194 (2017).

57 Tran, J. C. et al. Mapping intact protein isoforms in discovery mode using top-down proteomics. Nature 480, 254–258, doi:10.1038/nature10575 (2011).

58 Wang, H. et al. Role of histone H2A ubiquitination in Polycomb silencing. Nature 431, 873–878, doi:10.1038/nature02985 (2004).

59 Milani, G. et al. Low PKCalpha expression within the MRD-HR stratum defines a new subgroup of childhood T-ALL with very poor outcome. Oncotarget 5, 5234–5245, doi:10.18632/oncotarget.2062 (2014).

60 Serafin, V. et al. Phosphoproteomic analysis reveals hyperactivation of mTOR/STAT3 and LCK/Calcineurin axes in pediatric early T-cell precursor ALL. Leukemia 31, 1007–1011, doi:10.1038/leu.2017.13 (2017).

61 Benyoucef, A. et al. UTX inhibition as selective epigenetic therapy against TAL1-driven T-cell acute lymphoblastic leukemia. Genes & development 30, 508–521, doi:10.1101/gad.276790.115 (2016).

62 Hashizume, R. et al. Pharmacologic inhibition of histone demethylation as a therapy for pediatric brainstem glioma. Nature medicine 20, 1394–1396, doi:10.1038/nm.3716 (2014).

63 Tavana, O. et al. HAUSP deubiquitinates and stabilizes N-Myc in neuroblastoma. Nature medicine 22, 1180–1186, doi:10.1038/nm.4180 (2016).

64 Carra, G. et al. Therapeutic inhibition of USP7-PTEN network in chronic lymphocytic leukemia: a strategy to overcome TP53 mutated/deleted clones. Oncotarget, doi:10.18632/oncotarget.16348 (2017).

65 An, T. et al. USP7 inhibitor P5091 inhibits Wnt signaling and colorectal tumor growth. Biochem Pharmacol 131, 29–39, doi:10.1016/j.bcp.2017.02.011 (2017).

66 Malapelle, U. et al. USP7 inhibitors, downregulating CCDC6, sensitize lung neuroendocrine cancer cells to PARP-inhibitor drugs. Lung Cancer 107, 41–49, doi:10.1016/j.lungcan.2016.06.015 (2017).

67 Morra, F. et al. The combined effect of USP7 inhibitors and PARP inhibitors in hormone-sensitive and castration-resistant prostate cancer cells. Oncotarget, doi:10.18632/oncotarget.16463 (2017).

68 Huether, R. et al. The landscape of somatic mutations in epigenetic regulators across 1,000 paediatric cancer genomes. Nat Commun 5, 3630, doi:10.1038/ncomms4630 (2014).

69 Richter-Pechanska, P. et al. Identification of a genetically defined ultra-high-risk group in relapsed pediatric T-lymphoblastic leukemia. Blood Cancer J 7, e523, doi:10.1038/bcj.2017.3 (2017).

70 Van der Meulen, J. et al. The H3K27me3 demethylase UTX is a gender-specific tumor suppressor in T-cell acute lymphoblastic leukemia. Blood 125, 13–21, doi:10.1182/blood-2014-05-577270 (2015).

## Reference

1 Zheng, Y., Thomas, P. M. & Kelleher, N. L. Measurement of acetylation turnover at distinct lysines in human histones identifies long-lived acetylation sites. Nat Commun 4, 2203, doi:10.1038/ncomms3203 (2013).

2 Zheng, Y. et al. Total kinetic analysis reveals how combinatorial methylation patterns are established on lysines 27 and 36 of histone H3. Proceedings of the National Academy of Sciences of the United States of America 109, 13549–13554, doi:10.1073/pnas.1205707109 (2012).

3 Cotto-Rios, X. M.., Bekes, M., Chapman, J., Ueberheide, B. & Huang, T. T. Deubiquitinases as a signaling target of oxidative stress. Cell Rep 2, 1475–1484, doi:10.1016/j.celrep.2012.11.011 (2012).

4 Bolger, A. M., Lohse, M. & Usadel, B. Trimmomatic: a flexible trimmer for Illumina sequence data. Bioinformatics 30, 2114–2120, doi:10.1093/bioinformatics/btu170 (2014).

5 Langmead, B., Trapnell, C., Pop, M. & Salzberg, S. L. Ultrafast and memory-efficient alignment of short DNA sequences to the human genome. Genome Biol 10, R25, doi:10.1186/gb-2009-10-3-r25 (2009).

6 Lawrence, M. et al. Software for computing and annotating genomic ranges. PLoS Comput Biol 9, e1003118, doi:10.1371/journal.pcbi.1003118 (2013).

7 Zhang, Y. et al. Model-based analysis of ChIP-Seq (MACS). Genome Biol 9, R137, doi:10.1186/gb-2008-9-9-r137 (2008).

8 Heinz, S. et al. Simple combinations of lineage-determining transcription factors prime cis-regulatory elements required for macrophage and B cell identities. Mol Cell 38, 576–589, doi:10.1016/j.molcel.2010.05.004 (2010).

9 Haiminen, N., Mannila, H. & Terzi, E. Determining significance of pairwise co-occurrences of events in bursty sequences. BMC Bioinformatics 9, 336, doi:10.1186/1471-2105-9-336 (2008).

10 McLean, C. Y. et al. GREAT improves functional interpretation of cis-regulatory regions. Nat Biotechnol 28, 495–501, doi:10.1038/nbt.1630 (2010).

11 Shen, L., Shao, N., Liu, X. & Nestler, E. ngs.plot: Quick mining and visualization of next-generation sequencing data by integrating genomic databases. BMC Genomics 15, 284, doi:10.1186/1471-2164-15-284 (2014).

12 Saldanha, A. J. Java Treeview‐‐extensible visualization of microarray data. Bioinformatics 20, 3246–3248, doi:10.1093/bioinformatics/bth349 (2004).

13 Wang, L. et al. Ubiquitin-specific Protease-7 Inhibition Impairs Tip60-dependent Foxp3+ T-regulatory Cell Function and Promotes Antitumor Immunity. EBioMedicine 13, 99–112, doi:10.1016/j.ebiom.2016.10.018 (2016).

14 Ntziachristos, P. et al. Contrasting roles of histone 3 lysine 27 demethylases in acute lymphoblastic leukaemia. Nature 514, 513–517, doi:10.1038/nature13605 (2014).

15 Trapnell, C., Pachter, L. & Salzberg, S. L. TopHat: discovering splice junctions with RNA-Seq. Bioinformatics 25, 1105–1111, doi:10.1093/bioinformatics/btp120 (2009).

16 Anders, S., Pyl, P. T. & Huber, W. HTSeq‐‐a Python framework to work with high-throughput sequencing data. Bioinformatics 31, 166–169, doi:10.1093/bioinformatics/btu638 (2015).

17 Robinson, M. D., McCarthy, D. J. & Smyth, G. K. edgeR: a Bioconductor package for differential expression analysis of digital gene expression data. Bioinformatics 26, 139–140, doi: 10.1093/bioinformatics/btp616 (2010).

18 Loven, J. et al. Selective inhibition of tumor oncogenes by disruption of super-enhancers. Cell 153, 320–334, doi:10.1016/j.cell.2013.03.036 (2013).

19 Whyte, W. A. et al. Master transcription factors and mediator establish super-enhancers at key cell identity genes. Cell 153, 307–319, doi:10.1016/j.cell.2013.03.035 (2013).

20 Subramanian, A. et al. Gene set enrichment analysis: a knowledge-based approach for interpreting genome-wide expression profiles. Proc Natl Acad Sci USA 102, 15545–15550, doi:10.1073/pnas.0506580102 (2005).

21 Mootha, V. K. et al. PGC-1alpha-responsive genes involved in oxidative phosphorylation are coordinately downregulated in human diabetes. Nat Genet 34, 267–273, doi:10.1038/ng1180 (2003).

